# Widespread inhibitory projections from the cerebellar interposed nucleus

**DOI:** 10.1101/2020.12.31.425011

**Authors:** Elena N. Judd, Samantha M. Lewis, Abigail L. Person

## Abstract

The cerebellum consists of parallel parasagittal modules that contribute to diverse behaviors, spanning motor to cognitive. Recent work employing cell-type specific tracing has identified circumscribed output channels of the cerebellar nuclei that could confer tight functional specificity. These studies have largely focused on excitatory projections of the cerebellar nuclei, however, leaving open the question of whether inhibitory neurons also constitute multiple output modules. We mapped output and input patterns to intersectionally restricted cell types of the interposed and adjacent interstitial nuclei. In contrast to the widespread assumption of primarily excitatory outputs and restricted inferior olive-targeting inhibitory output, we found that inhibitory neurons from this region ramified widely within the brainstem, targeting both motor- and sensory-related nuclei, distinct from excitatory output targets. Despite differences in output targeting, monosynaptic rabies tracing revealed largely shared afferents to both cell classes. We discuss the potential novel functional roles for inhibitory outputs in the context of cerebellar theory.

## Introduction

The cerebellum plays a critical role in refining motor control through learning. The cerebellar nuclei (CbN), which constitute the major outputs of the cerebellum, are proposed to relay predictive computations of the cerebellar cortex and store well-learned patterns, placing them in a central position to implement cerebellar control (Eccles et al., 1974; Ohyama et al., 2003; Chan-Palay, 1977). The CbN are a collection of nuclei that house diverse neuronal subtypes that differ in their targets. Recent studies have greatly expanded our understanding of this diversity, using approaches such as genomic profiling and projection specific tracing (Bagnall et al., 2009; Low et al., 2018; Fujita et al., 2020; Kebschull et al., 2020; Uusisaari & Knöpfel, 2010, 2011; Uusisaari et al., 2007; Husson et al., 2014; Ankri et al., 2015; Canto et al., 2016). Through these studies, we know that multiple diverse output channels intermingle (Fujita et al., 2020; Low et al., 2018; Sathyamurthy et al., 2020), widespread collateralization is common, and genetic diversity of excitatory projection neurons varies systematically along the medio-lateral extent of the cerebellar nuclei which encompasses the medial (fastigial), interposed, lateral (dentate), interstitial, and vestibular nuclei (Kebschull et al., 2020).

The mouse cerebellar interposed nucleus has received recent attention at the anatomical and functional levels with studies identifying specific projection patterns and functional roles for neuronal subtypes within the structure. Interposed excitatory neurons project to a variety of motor-related spinal cord and brainstem targets, as well as collateralize to motor thalamus (Low et al., 2018; Sathyamurthy et al., 2020; Kebschull et al., 2020). Ablation of a subset of anterior interposed (IntA) glutamatergic cells that express Urocortin3, for example, disrupts accurate limb positioning and timing during a reach to grasp task and locomotion (Low et al., 2018). Chemogenetic silencing of excitatory neurons that project ipsilaterally to the cervical spinal cord also impaired reach success in mice (Sathyamurthy et al., 2020). Moreover, closed-loop manipulation of IntA disrupts reach endpoint in real time (Becker & Person, 2019). The interposed nucleus also mediates conditioned eyelid responses, sculpts reach and gait kinematics, and is responsive to tactile stimulation (Darmohray et al., 2019; ten Brinke et al., 2017; Rowland and Jaeger, 2005). How anatomical organization of the structure confers such functions is an open question.

Functional consequences of cell type specific manipulations have not been limited to excitatory neurons. Ablation of inhibitory nucleo-olivary cells demarcated with Sox14 expression also resulted in motor coordination deficits (Prekop et al., 2018). These cells were traced from the lateral nucleus and suggested to project solely to the inferior olive (IO), consistent with conclusions from experiments using dual labeling methods (Ruigrok and Teune, 2014). Nevertheless, older reports of inhibitory projections from the cerebellar nuclei that target regions other than the IO raise the question of whether inhibitory outputs might also play a role in regulating brainstem nuclei outside the olivocerebellar system. Combined immunostaining with horseradish peroxidase tracing from the basilar pontine nuclei (i.e. pontine gray) in rats and cats showed GABA immunopositive neurons in the lateral nucleus (Aas and Brodal, 1989; Border et al., 1986), although the literature is inconsistent (Schwarz & Schmitz, 1997). Glycinergic output projections from the medial nucleus (fastigial) inhibitory output population includes large glycinergic neurons that project to ipsilateral brainstem targets outside the IO (Bagnall et al., 2009), unlike its Gad2-expressing neurons which exclusively target the IO (Fujita et al., 2020). In aggregate, these various observations indicate that better understanding of whether the interposed nucleus houses inhibitory output neurons that project to targets outside IO is an important open question.

Here we use a range of viral tracing methods to isolate and map projections from and to inhibitory and excitatory neurons of the intermediate cerebellar nuclear groups, defined through intersectional labeling methods using single or multiple recombinases coupled with pathway-specific labeling (Fenno et al., 2014). This method permitted analysis of collateralization more specific than traditional dual-retrograde labeling strategies since it leverages genetic specification and projection specificity and permits entire axonal fields to be traced. We elucidate the projection “fingerprints” of genetic- and projection-defined cell groups. Surprisingly, we observed widespread inhibitory outputs, comprised in part of putative collaterals of IO-projecting neurons, that target both ipsilateral and contralateral brainstem and midbrain structures. Monosynaptic rabies transsynaptic tracing (Kim et al., 2016; Wickersham et al., 2010) restricted to excitatory premotor neuron populations through the selective expression of Cre recombinase under the Vglut2 promoter (Gong et al., 2007) and inhibitory neurons through Cre expression controlled under the Vgat promoter revealed reproducible patterns of presynaptic inputs largely shared across cell types. Together these experiments provide new insight into input/output organization of the intermediate cerebellum, suggest potential functional diversity of parallel channels, and provide anatomical targets for functional studies aimed at evaluating these putative roles.

## Results

### Anterograde tracing of Int-Vgat neurons

To determine projection patterns of inhibitory neurons of the interposed nucleus, we stereotaxically injected AAV2.EF1a.DIO.YFP into Vgat-Cre transgenic mice, “Int-Vgat”, (N = 5, Fig. 1A). We mapped and scored the extent and density of terminal varicosities on a 4 point scale and recorded injection sites, plotted for all experiments (Fig. S1; See Methods, Projection quantification).

**Figure 1.**
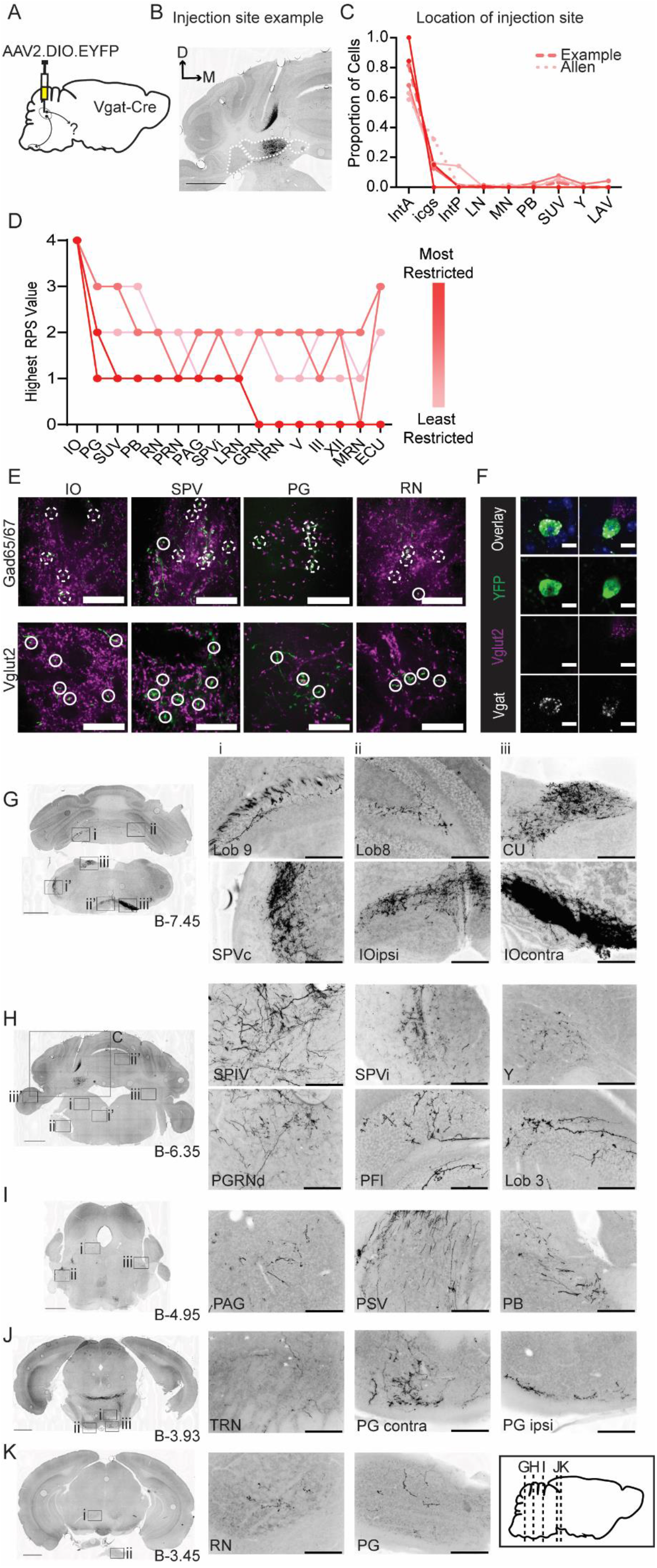
Anterograde tracing of Int-Vgat neurons. (A) Schematic of injection scheme. (B) Example injection site of AAV2-EF1a-DIO-eYFP. The three main CbN are outlined in white (*Lateral Nucleus (LN), Interposed (IN), and Medial Nucleus (MN)* from left to right). Images oriented so that the dorsal ventral axis runs up/ down and the medial/ lateral axis runs right/ left; right of midline is contralateral. (C) Location of labeled cells by injection into Int of Vgat-Cre mice. Specimens are color coded by proportion of cells labeled in anterior interposed (IntA) where the highest proportion corresponds to darkest color. (D) Mapping of terminal fields based on restriction of injection site to IntA. The highest unilateral RPS in each region is plotted for all specimens included in analysis. (E) YFP-positive terminals (green) in *inferior olive (IO), spinal trigeminal nuclei (SPVc), pontine grey (PG), and red nucleus (RN)* are stained for antibodies against Gad65/67 (top) and Vglut2 (bottom; magenta). Dashed circles indicate colocalized terminals while solid lines indicate a lack of colocalization observed in the two channels. Scale bars = 20 µms. (F) Example cells from *in situ* hybridization showing clear overlap with an mRNA probe against SLC32a1 (Vgat) and non overlap with an mRNA probe against SLC17a6 (Vglut2). Scale bars = 10 µms. (G) Projection targets in caudal cerebellum and brainstem (B-7.45). Boxes expanded in i-iii (top) or i-iii’ (bottom). (H) Projection targets within the intermediate cerebellum (B-6.35). Injection site depicted in C. (I) Projection targets within rostral brainstem (B-4.95). (J) Projection targets in the caudal midbrain (B-3.93). (K) Projection targets to the rostral midbrain (B-3.93). Scale bars (C, G-K) = 1 mm and (i-iii) 200 µms. The inset (black border) depicts the location of coronal sections shown in G-K along a parasagittal axis. *Cuneate nucleus (CU), gigantocellular reticular nucleus (GRN), hypoglossal nucleus (XII), intermediate reticular nucleus (IRN), interstitial cell groups (icgs), lateral reticular nucleus (LRN), lateral vestibular nucleus (LAV), midbrain reticular nucleus (MRN), motor nucleus of the trigeminal (V), nucleus prepositus (PRP), Nucleus Y (Y), oculomotor nucleus (III), parabrachial (PB), paraflocculus (PFl), paragigantocellualr reticular nucleus (PGRN), periaqueductal grey (PAG), principle sensory nucleus of the trigeminal (PSV), pontine reticular nucleus (PRN), posterior interposed (IntP), spinal trigeminal nucleus, caudal/ interpolar subdivision (SPVc/i), spinal vestibular nucleus (SPIV), superior vestibular nucleus (SUV), tegmental reticular nucleus (TRN)*.

As expected, injections labeled neurons that densely innervated the contralateral dorsal accessory inferior olive (IO; Fig. 1D,E,G), with less dense but consistent innervation of ipsilateral IO (Ruigrok and Voogd, 1990; Balaban & Beryozkin, 1994; Fredette & Mugnaini, 1991; Prekop et al., 2018; Ruigrok & Voogd, 1990, 2000; Want et al., 1989). Surprisingly these injections also consistently labeled terminal fields outside IO, within the brainstem, even when injection sites were completely restricted to the anterior interposed nucleus (Fig. S5). Viral expression of Int-Vgat neurons labeled axonal varicosities which were immunopositive for probes against Gad65/67, but never Vglut2, consistent with a GABAergic phenotype for these projections (Fig. 1E, S2 analyzed in the inferior olive, interpolar spinal trigeminal nucleus (SPVi), pontine gray (PG), red nucleus (RN) and vestibular nuclei). *In situ* hybridization revealed that 98% of virally labeled cells co-expressed the Vgat marker *Slc32a1* (230/234 cells from 2 mice), while 4/234 cells overlapped the glutamatergic marker *Slc17a6* (Fig. 1F, S3, Table S1). A Gad1-Cre driver line (Higo et al., 2007) was tested but not used owing to non-specific label (Fig. S4; See Methods; Table S1).

Most Int-Vgat injections included both interposed and interstitial cell groups slightly ventral to the interposed nucleus, plotted in Fig.1B, color coded for the percentage of the injection site contained within IntA. Although injection site spillover into interstitial cell groups (Sugihara and Shinoda, 2007) was common, injection site spillover into the main vestibular groups ventral to the 4^th^ ventricle was minimal to absent. Following these injections, terminal label within the brainstem was extensive, and invariably also included beaded varicosities within the cerebellar cortex characteristic of the inhibitory nucleocortical pathway (Ankri et al, 2015). Modestly dense but spatially extensive terminal fields ramified in the posterior medulla along the anterior-posterior axis (Fig. 1, Fig. 2D). Among sensory brainstem structures, terminal fields ramified within the ipsilateral external cuneate nucleus (ECU), cuneate nucleus (CU), nucleus of the solitary tract (NTS), spinal trigeminal nucleus (SPVi), especially the lateral edge, parabrachial nuclei (PB), and principal sensory nuclei of the trigeminal (PSV), and all vestibular nuclei. Int-Vgat axons extended through the pontine reticular nuclei (PRN) to innervate the tegmental reticular nuclei (TRN; commonly abbreviated NRTP) and the pontine gray (PG; i.e. basilar pontine nuclei; Fig. 1D, J), which are themselves major sources of cerebellar mossy fibers. Int-Vgat neurons also innervated the medial magnocellular red nucleus (RN) (Fig. 1K) bilaterally. Rarely, Int-Vgat axons progressed to the caudal diencephalon, very sparsely targeting the ipsilateral zona incerta ZI in 2/6 mice (Table S2). Axonal varicosities were vanishingly sparse or non-existent within the spinal cord following Int-Vgat injections (data not shown).

**Figure 2.**
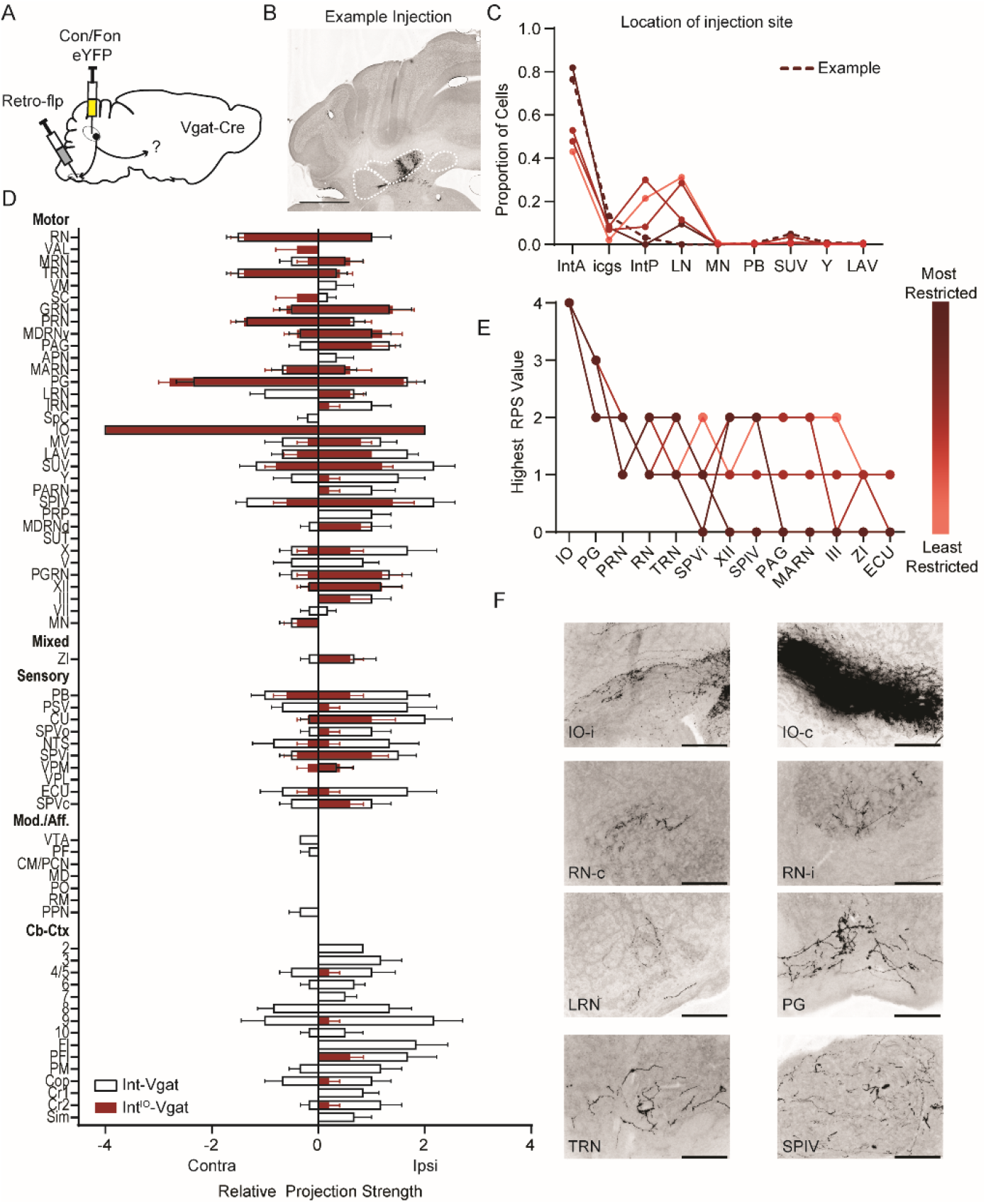
Intersectional labeling of IO-projecting Int-Vgat neurons (Int^IO^-Vgat) and comparison with Int-Vgat. (A) Schematic of experiment. (B) Example injection site of AAV8.hSyn.Con/Fon.hChR2.EYFP in a Vgat-Cre mouse. The three main CbN are outlined in white (*Lateral Nucleus (LN), Interposed (IN), and Medial Nucleus (MN)* from left to right). Images oriented as in Fig. 1. Scale bar = 1 mm. (C) Location of labeled cells by injection of Retro-Flp to the contralateral *inferior olive (IO)* and Con/Fon-YFP into Int of Vgat-Cre mice. Specimens are color coded by proportion of cells labeled in *anterior interposed (IntA)* where the highest proportion corresponds to darkest color. (D) Graphical representation of average projection strength in all targeted regions for Int^IO^-Vgat (n = 5; maroon) and Int-Vgat (n = 6; white) mice. See Table 1 for full list of abbreviations. (E) Mapping of terminal fields based on restriction of injection site to IntA. The highest unilateral RPS in each region is plotted for all specimens included in analysis. (F) Example terminal fields within the *inferior olive (IO) and red nucleus (RN) bilaterally, lateral reticular nucleus (LRN), pontine grey (PG), tegmental reticular nucleus (TRN), and spinal vestibular nucleus (SPIV).* Scale bars = 200 µms. *External cuneate nucleus (ECU), hypoglossal nucleus (XII), interstitial cell groups (icgs), lateral vestibular nucleus (LAV), magnocellular reticular nucleus (MARN), Nucleus Y (Y), oculomotor nucleus (III), parabrachial (PB), periaqueductal grey (PAG), pontine reticular nucleus (PRN), posterior interposed (IntP), spinal trigeminal nuclei, interpolar (SPVi), spinal vestibular nucleus (SPIV), superior vestibular nucleus (SUV), zona incerta (ZI)*.

Beaded nucleocortical fibers from Int-Vgat injections were reliably labeled if the injection site included interstitial cell groups (Fig. 1 G,H, 2D; Ankri et al., 2015). Int-Vgat neurons targeted all cerebellar lobules, even extending contralaterally. Several specimens showed minor label of inhibitory cells in the ventral Cb-Ctx just dorsal to IntA, but nucleocortical terminals that were included in the projection analysis were not located in the same topographical area.

Some targets noted were sensitive to injection site restriction (Fig. 1D). However, labeling of varicosities outside IO was not attributable solely to injection site leakage outside Int. The smallest Int-Vgat injection, contained entirely within IntA, labeled fine caliber axons that ramified within the ipsilateral superior and spinal vestibular nuclei (Fig. S5). Labelled fibers coursed in the superior cerebellar peduncle, decussating at the level of the pontine nuclei (-4 mm Bregma). As they coursed ventrally, they produced numerous varicosities in the pontine nuclei, specifically the tegmental reticular nucleus and pontine gray, before turning caudally, labeling dense terminals fields in the contralateral IO (DAO) and modestly dense fields in the ipsilateral IO. Very sparse varicosities were also noted in the parabrachial nucleus and magnocellular red nucleus. Despite the presence of these terminal fields, no nucleocortical fibers were seen following the most restricted Int-Vgat injection, suggesting these may originate from interstitial cell groups. To summarize, Int-Vgat injections labeled fibers that innervated numerous brainstem nuclei outside IO, even following highly restricted injections.

### Projection-specific Int-Vgat neuron tracing

The terminals observed in brainstem and midbrain from Int-Vgat labeling suggested the existence of inhibitory channels from the intermediate cerebellum beyond those targeting the IO. Next, to restrict label to genetic- and projection-specific Int neurons (Fenno et al., 2014), we used a two-recombinase-dependent reporter virus (AAV8.hsyn.Con/Fon.eYFP) injected into Int in conjunction with Flp recombinase retrogradely introduced via the contralateral IO with AAVretro-EF1a-Flp (Fig. 2A; N = 5). The fluorescent reporter will only express in the presence of both Cre and Flp recombinases. This Cre-on Flp-on approach, termed “Con/Fon”, was used to isolate IO-projecting Int-Vgat neurons. Specificity was determined via injections in wildtype C57/Bl6 mice (N=2) and off-target injections in Cre mice (N=3), which did not yield YFP positive neurons in the cerebellar nuclei (Fig. S6).

Int^IO^-Vgat neurons had more restricted terminations than most direct Int-Vgat injections. Varicosities were consistently observed in dorsal PG, PRN, TRN, IO and the vestibular complex. Less consistent and sparser label occurred in other brainstem nuclei (Fig. 2). These data suggest that IO-projecting cells collateralize to a subset of targets relative to the constellation of regions targeted by all Int-Vgat neurons, typically excluding nucleocortical projections, modulatory/affective regions, and sensory nuclei.

### Anterograde tracing from excitatory output neurons

To compare Int-Vgat projections more directly to excitatory outputs, we injected Int of Ntsr1-Cre mice with AAV1.CAG.flex.GFP (N=2) or AAV2.DIO.EF1a.eYFP (N=3) (Fig. 3). Int-Ntsr1 terminal varicosities consistently colocalized with Vglut2 immunolabel, but never Vgat, consistent with a glutamatergic phenotype of Ntsr1 output neurons (Fig. 3E, S7), and somata overlapped predominantly with the glutamatergic marker *Slc17a6* (Fig. 3F; S3). Dense and consistent terminal varicosities labeled by Int-Ntsr1 neurons occurred in patches within the caudal medulla, midbrain, and thalamus, which are known targets of Vglut2-Cre and Ucn3-Cre neurons (Fig. 3D, G-K, Low et al., 2018; Kebschull et al., 2020; Sathyamurthy et al., 2020). Varicosities filled the ipsilateral parvicellular reticular nucleus PARN (commonly abbreviated PCRt) which extended rostrally to blend into the spinal nucleus of the trigeminal (SPV), known forelimb control structures (Esposito et al., 2014), and ipsilateral terminals ramified in the motor nucleus of the trigeminal (V). Bilateral patches of terminals were seen in the lateral reticular nucleus (LRN) and all four subdivisions of the vestibular nuclei. At the level of the decussation of the superior cerebellar peduncle, axons turned ventrally and produced dense Vglut2-positive varicosities in the TRN (commonly abbreviated NRTP) and sparsely in PG (Cicirata et al. 2005; Schwarz and Schmitz 1997). Axons also ramified within the magnocellular RN and the deep layers of the SC. Diencephalic projections were densely targeted to thalamic nuclei and more sparsely targeted to ZI. All specimens exhibited dense terminal fields in the ventromedial (VM) and anterior ventrolateral (VAL) nuclei of the thalamus (Teune et al 2000; Aumann et al., 1994; Houck & Person, 2015; Kalil, 1981; Low et al., 2018; Stanton, 1980). Additionally, we observed terminals in intralaminar thalamic structures including: centromedial (CM), paracentral (PCN), mediodorsal (MD), parafascicular (PF), ventral posterior (VP), and posterior (PO) nuclei (Teune et al., 2000; Chen et al., 2014; Dumas et al., 2019). Int-Ntsr1 neurons formed nucleocortical mossy fibers in multiple lobules across the cortex (Fig. 3G-I; Gao et al., 2016; Houck & Person, 2015; Tolbert et al., 1978; Low et al., 2018; Sathyamurthy et al., 2020).

**Figure 3.**
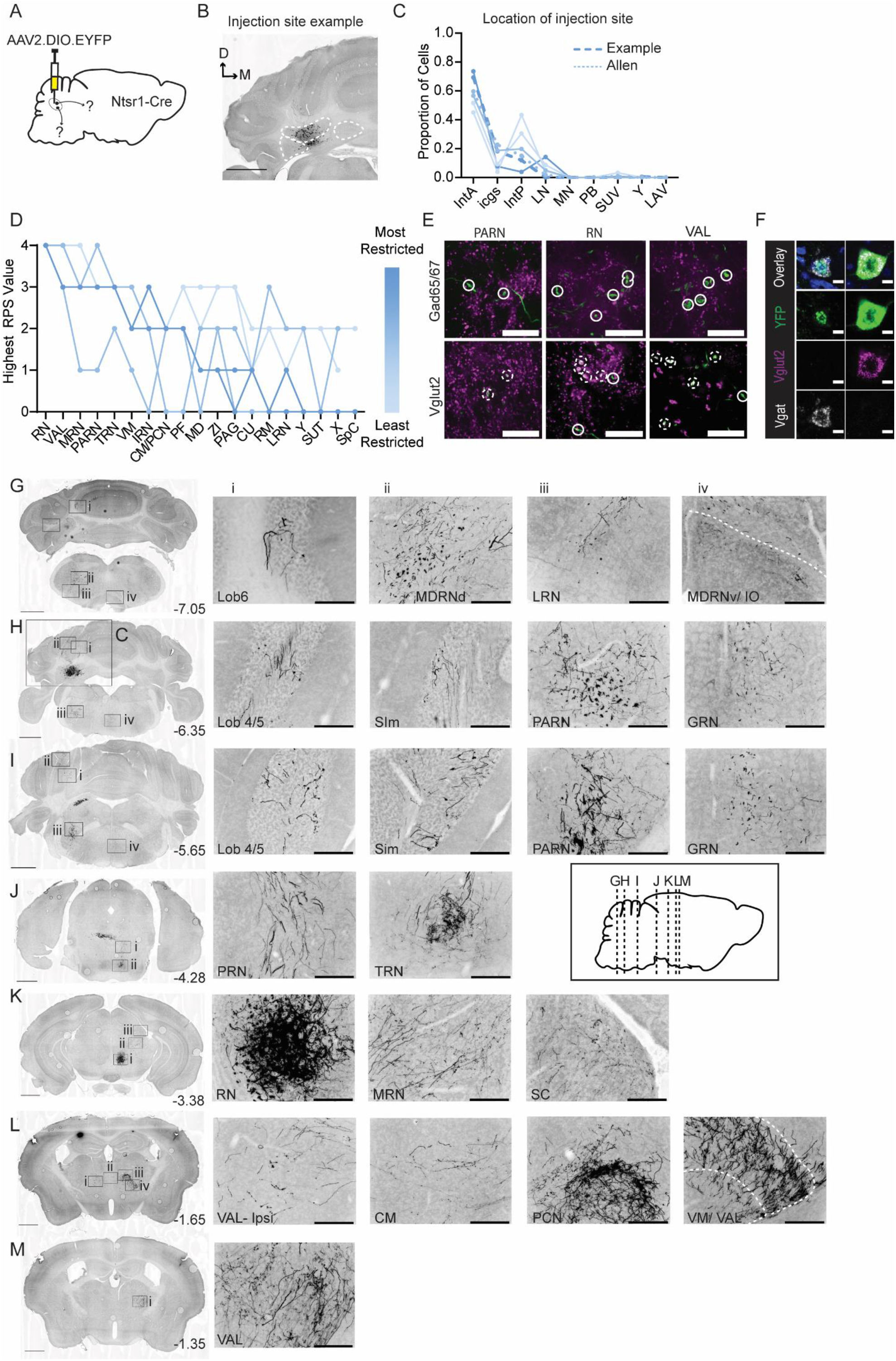
Anterograde tracing of Int-Ntsr1 neurons. (A) Schematic representation of injection scheme. (B) Example injection site of AAV2-EF1a-DIO-eYFP. The three main CbN are outlined in white *(Lateral Nucleus (LN), Interposed (IN), and Medial Nucleus (MN)* from left to right). Images oriented as in Fig.1. (C) Distribution of labeled cells by injection into Int of Ntsr1-Cre mice. Specimens are color coded by proportion of cells labeled in *anterior interposed (IntA)* where the highest proportion corresponds to darkest color. (D) Mapping of terminal fields based on restriction of injection site to IntA. The highest unilateral RPS in each region is plotted for all specimens included in analysis. (E) YFP-positive terminals (green) in *pavicellular reticular nucleus (PARN), red nucleus (RN), and ventral anterior-lateral complex of the thalamus (VAL)* are stained for antibodies against Gad65/67 (top; magenta) and Vglut2 (bottom; magenta). Dashed circles indicate colocalized terminals while solid lines indicate a lack of colocalization observed in the two channels. Scale bars = 20 µms. (F) Example cells from *in situ* hybridization showing overlap with both an mRNA probe against SLC32a1 (Vgat) and SLC17a6 (Vglut2). Scale bars = 10 µms. (G) Projection targets in caudal cerebellum and brainstem (B-7.05). Boxes expanded in i-iv. (H) Projection targets within the intermediate cerebellum (B-6.35). Injection site depicted in C. (I) Projection targets within and ventral to the anterior cerebellum (B-5.65). (J) Projection targets to pontine nuclei (B-4.25). (K) Projection targets in the rostral midbrain (B-3.38). Note the dense terminals in RN. (L) Projection targets to the caudal thalamus (B-1.65). (M) Projection targets to the rostral thalamus (B-1.35). Scale bars (C, G-M) = 1 mm and (i-iv) 200 µms. The inset (black border) depicts the location of coronal sections shown in G-M along a parasagittal axis. *Centromedial nucleus of the thalamus (CM), cuneate nucleus (CU), gigantocellular reticular nucleus (GRN), inferior olive (IO), intermediate reticular nucleus (IRN), interstitial cell groups (icgs), lateral reticular nucleus (LRN), lateral vestibular nucleus (LAV), mediodorsal nucleus of the thalamus (MD), medullary reticular nucleus, dorsal/ ventral subdivision (MDRNd/v), midbrain reticular nucleus (MRN), nucleus raphe magnus (RM), nucleus X (X), nucleus Y (Y), parabrachial (PB), paracentral nucleus of the thalamus (PCN), parafascicular nucleus (PF), periaqueductal grey (PAG), pontine reticular nucleus (PRN), posterior interposed (IntP), simplex lobule (Sim), superior colliculus (SC), superior vestibular nucleus (SUV), supratrigeminal nucleus (SUT), spinal cord (SpC), tegmental reticular nucleus (TRN), ventromedial nucleus (VM), zona incerta (ZI)*.

Beyond the major targets described above, Int-Ntsr1 projected sparsely to a variety of other regions. In 3 of 5 animals, we observed a small patch of terminals within the contralateral dorsal subnucleus of IO that were positive for Vglut2 (Fig. 3G, S7). Near the dense terminal field within the contralateral RNm, fine caliber axons bearing varicosities spilled over into the ventral tegmental area, VTA (Figure S8; Carta et al., 2019; Teune et al., 2000) and extended dorsally through the contralateral midbrain/mesencephalic reticular nucleus (MRN; Ferreira-Pinto et al., 2021) to innervate the caudal anterior pretectal nucleus (APN) anterior ventrolateral periaqueductal grey (PAG) (Vaaga et al., 2020; Sugimoto et al., 1982; Gayer & Faull, 1988; Low et al., 2018; Teune et al., 2000). To summarize, Int-Ntsr1 neurons targeted regions well known to receive excitatory input from the interposed nucleus, as well as a previously unappreciated vGlut2+ afferent to the IO.

### Projection-specific Int-Ntsr1 neuron tracing

We next used the Con/Fon intersectional approach described above to restrict labeling to RN-projecting Ntsr1-Cre neurons (Int^RN^-Ntsr1, Fig. 4; N=4), asking whether projection-specific labeling recapitulated data from direct label of Int-Ntsr1 cells, as would be expected if RN projecting neurons collateralize to other targets. The projection pattern of Int^RN^-Ntsr1 was almost identical to the pattern observed in Int-Ntsr1 injections, with a few notable exceptions. Namely, only Int-Ntsr1 neurons projected to lobule 8, anterior pretectal nucleus (APN), IO, and pedunculopontine nuclei (PPN). Terminal fields in the contralateral thalamus, especially VAL, VM, and CM/ PCN as well as layers 7/8 of the contralateral cervical spinal cord (2/3 specimens with spinal cords available) support the observation in Sathyamurthy et al. (2020) that contralaterally projecting cerebellospinal neurons collateralize to both RN and thalamus. We conclude that it is likely that Int-Ntsr1 neurons reliably project to RN and collateralize to a restricted collection of other targets, although these data do not distinguish between broad vs restricted collateralization of Int^RN^-Ntsr1 neurons.

**Figure 4.**
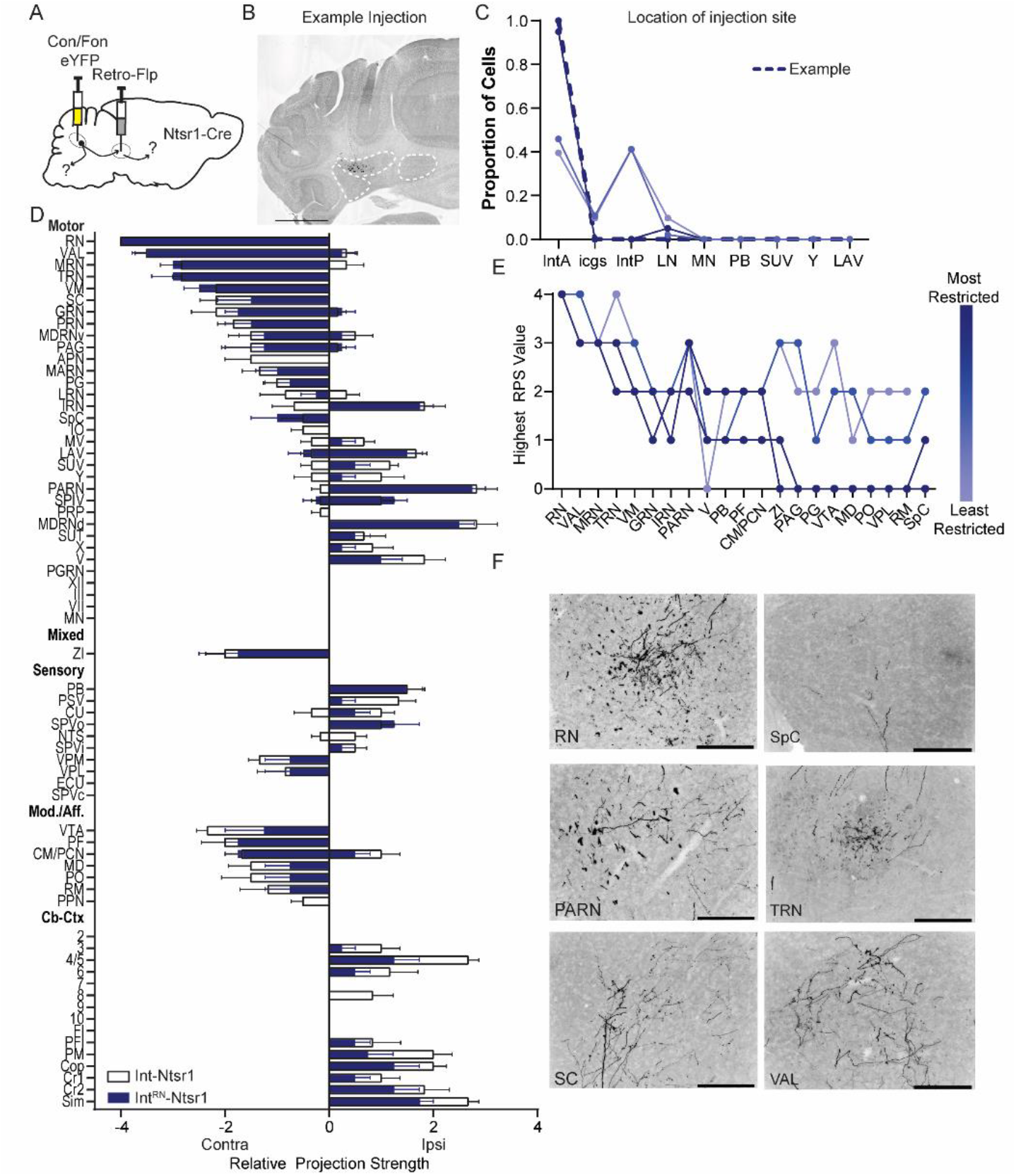
Intersectional labeling of RN-projecting Int-Ntsr1 neurons (Int^RN^-Ntsr1). (A) Schematic of experiment. (B) Example injection site of AAV8.hSyn.Con/Fon.hChR2.EYFP in an Ntsr1-Cre mouse. The three main CbN are outlined in white (*Lateral Nucleus (LN), Interposed (IN), and Medial Nucleus (MN) from left to right*). Images oriented so right of midline is contralateral. Scale bar = 1 mm. (C) Location of labeled cells by injection of Retro-Flp to the contralateral red nucleus (RN) and Con/Fon-YFP into Int of Ntsr1-Cre mice. Specimens are color coded by proportion of cells labeled in anterior interposed (IntA) where the highest proportion corresponds to darkest color. (D) Graphical representation of average projection strength in all targeted regions for Int^RN^-Ntsr1 (n = 4; navy) and Int-Ntsr1 (n = 6; white) mice. See Table 1 for a full list of abbreviations. (E) Mapping of terminal fields based on restriction of injection site to IntA. The highest unilateral RPS in each region is plotted for all specimens included in analysis. (F) Example terminal fields within the *red nucleus (RN), spinal cord (SpC), parvicellular reticular nucleus (PARN), tegmental reticular nucleus (TRN), superior colliculus (SC), and ventral anterior-lateral complex of the thalamus (VAL).* Scale bars = 200 µms. *Centromedial nucleus of the thalamus (CM), gigantocellular reticular nucleus (GRN), inferior olive (IO), intermediate reticular nucleus (IRN), interstitial cell groups (icgs), lateral reticular nucleus (LRN), lateral vestibular nucleus (LAV), mediodorsal nucleus of the thalamus (MD), midbrain reticular nucleus (MRN), motor nucleus of the trigeminal (V), nucleus raphe magnus (RM), nucleus X (X), nucleus Y (Y), parabrachial (PB), paracentral nucleus of the thalamus (PCN), parafascicular nucleus (PF), periaqueductal grey (PAG), pontine grey (PG), pontine reticular nucleus (PRN), posterior complex of the thalamus (PO), posterior interposed (IntP), superior vestibular nucleus (SUV), supratrigeminal nucleus (SUT), tegmental reticular nucleus (TRN), ventral tegmental area (VTA), ventromedial nucleus (VM), ventral posterolateral nucleus of the thalamus (VPL), zona incerta (ZI)*.

### Projections of IntA^RN^ neurons traced with AAVretro-Cre

As described above, we noted that both Int-Vgat and Int-Ntsr1 labeled varicosities within RN. This presented a target we could exploit to test whether Int neurons collateralize to both RN and IO independent of genetic Cre label. We retrogradely expressed Cre in RN-projecting neurons, injecting AAV2retro.Cre into RN and a flexed reporter virus into Int (AAV1.CAG.flex.GFP/ RFP) of wild type C57/Bl6 mice (Fig S9; N = 4). Following these injections, we observed label in both IO and RN contralateral to the Int injection (Fig. S9; Table 1, S2). We also observed terminals in other locations consistently targeted by either Int-Ntsr1 (MRN, VAL, VPM, VM, PF, MD, PO, SC, ZI) or Int-Vgat (Lob 9, IO, lateral SPV, ipsilateral PRN, and ECU). Following these injections, terminal varicosities in IO, and subsets in TRN and PG expressed Gad65/67 while varicosities in SPV, RN, PG, VAL and TRN were positive for Vglut2 (Fig. S10; N=2). We conclude that retrograde uptake of Cre from synaptic terminals in RN results in reporter expression of both glutamatergic and GABAergic neurons in Int that both project to RN.

**Table 1.**
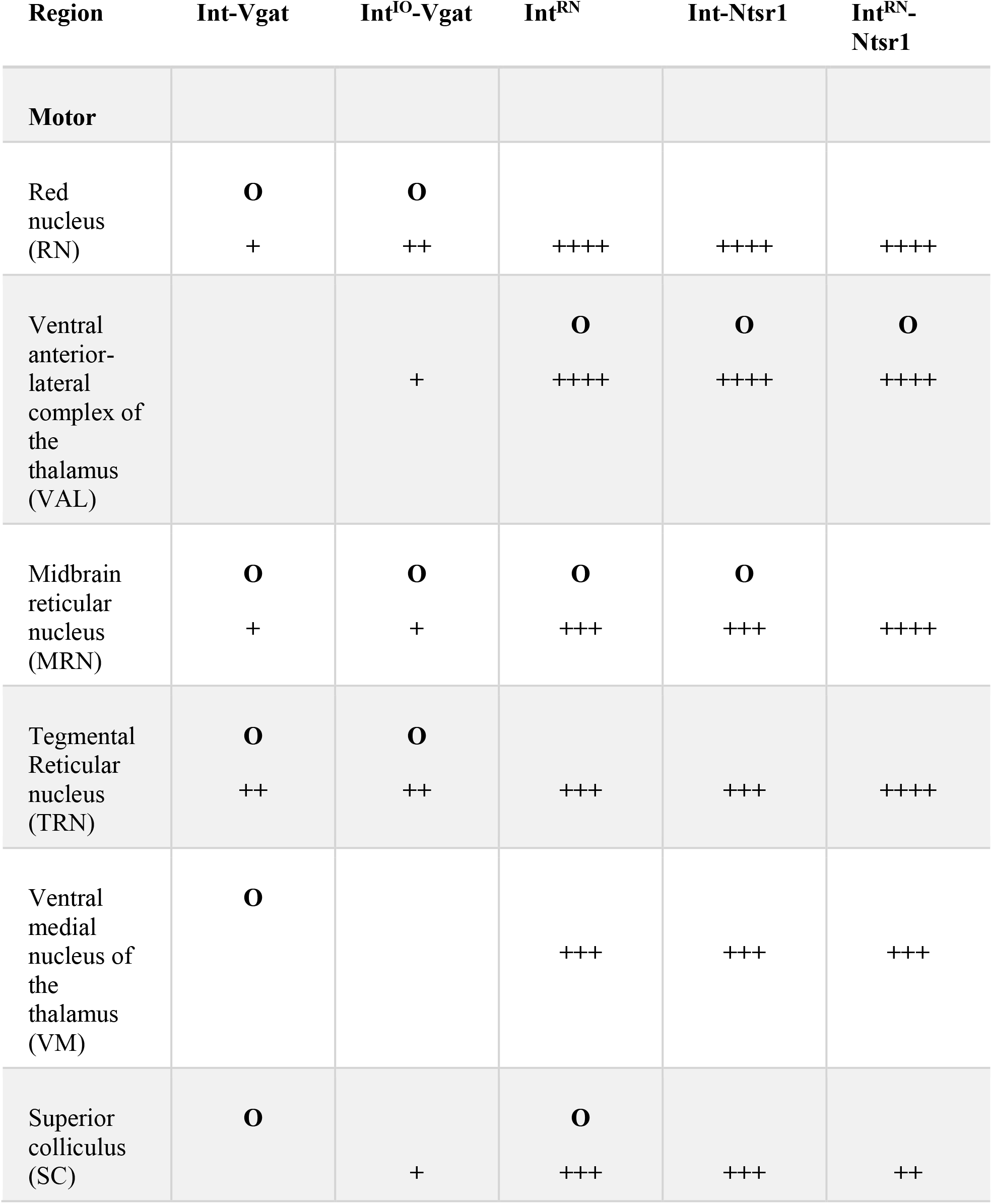
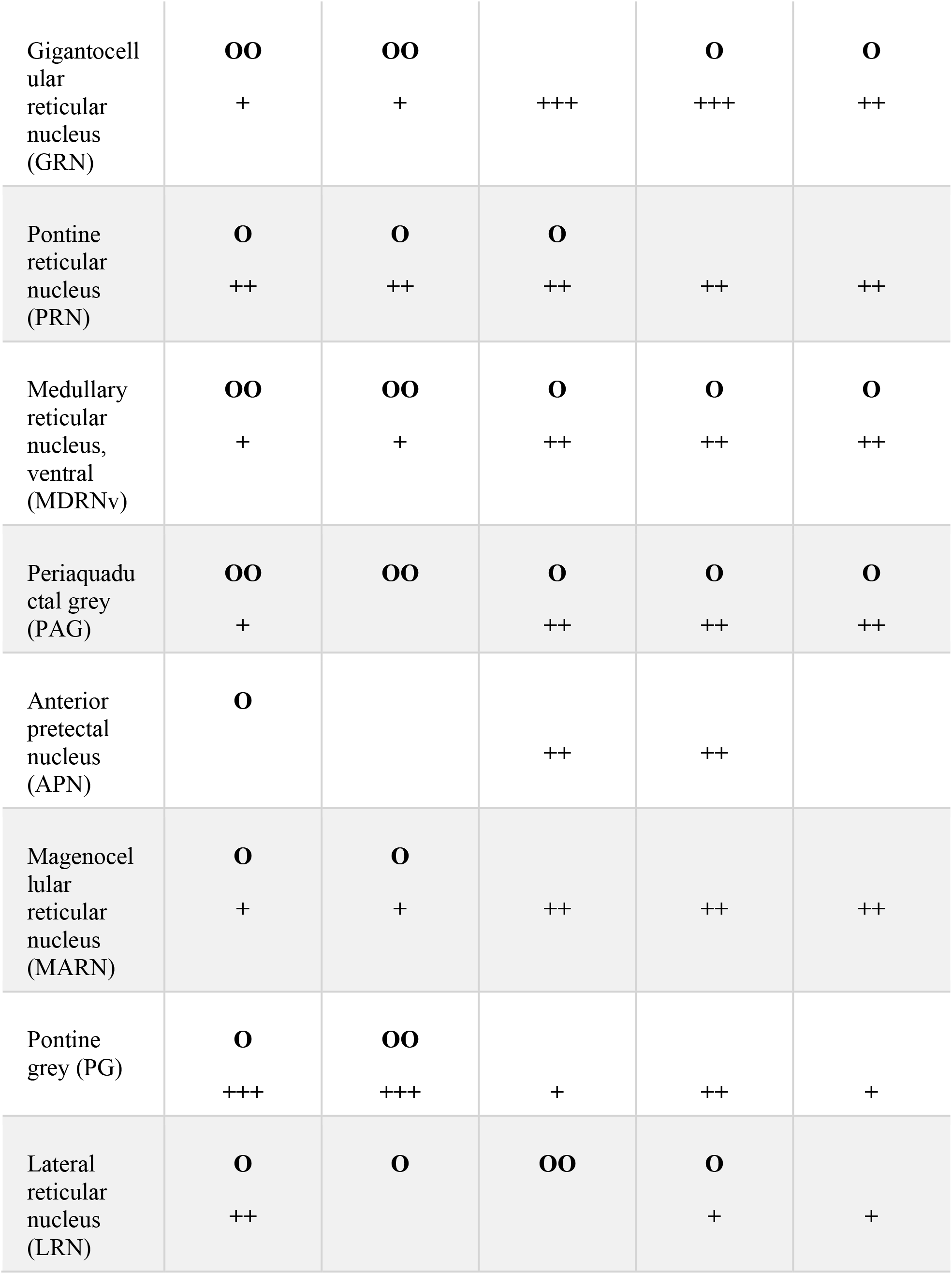
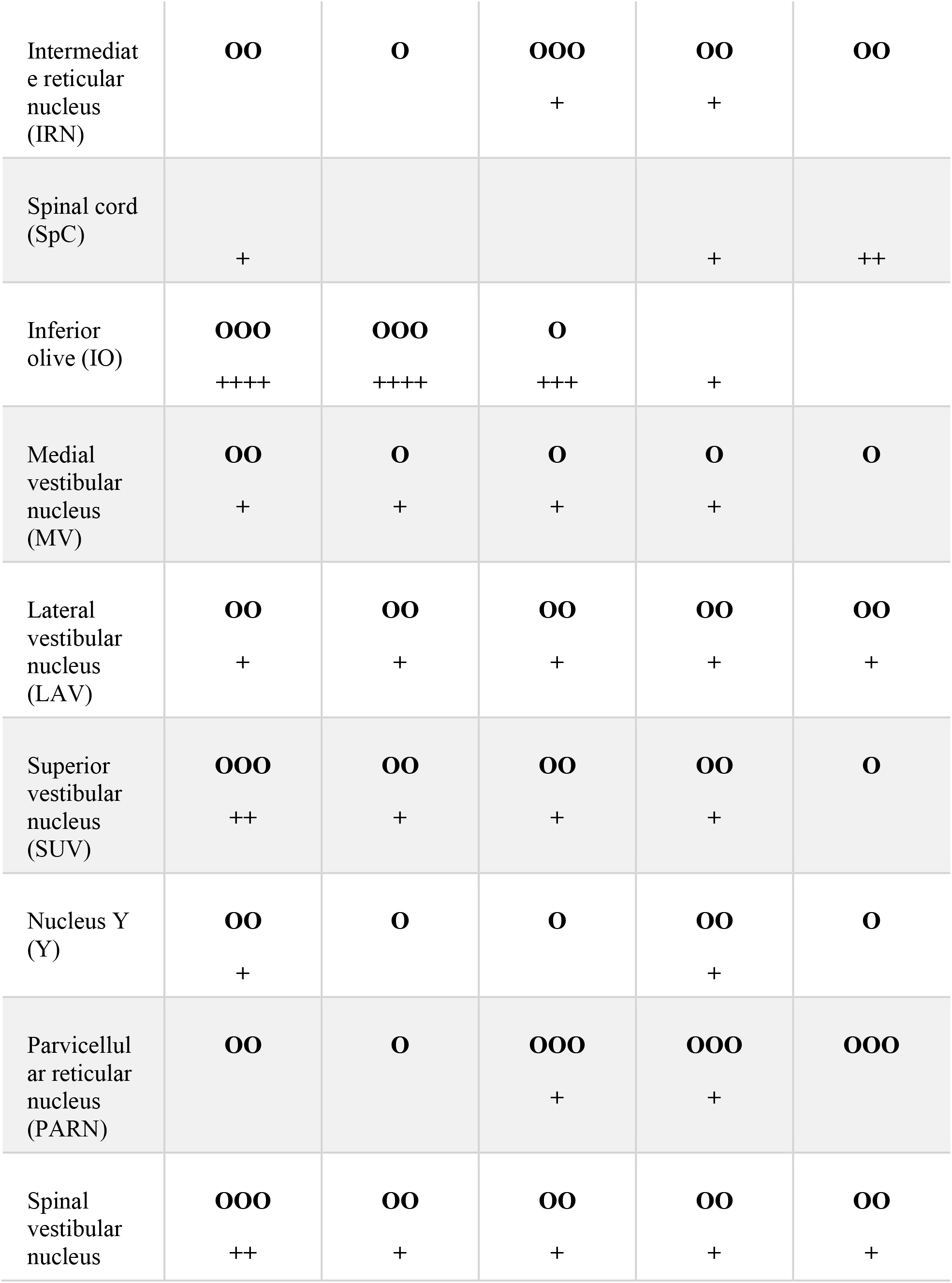
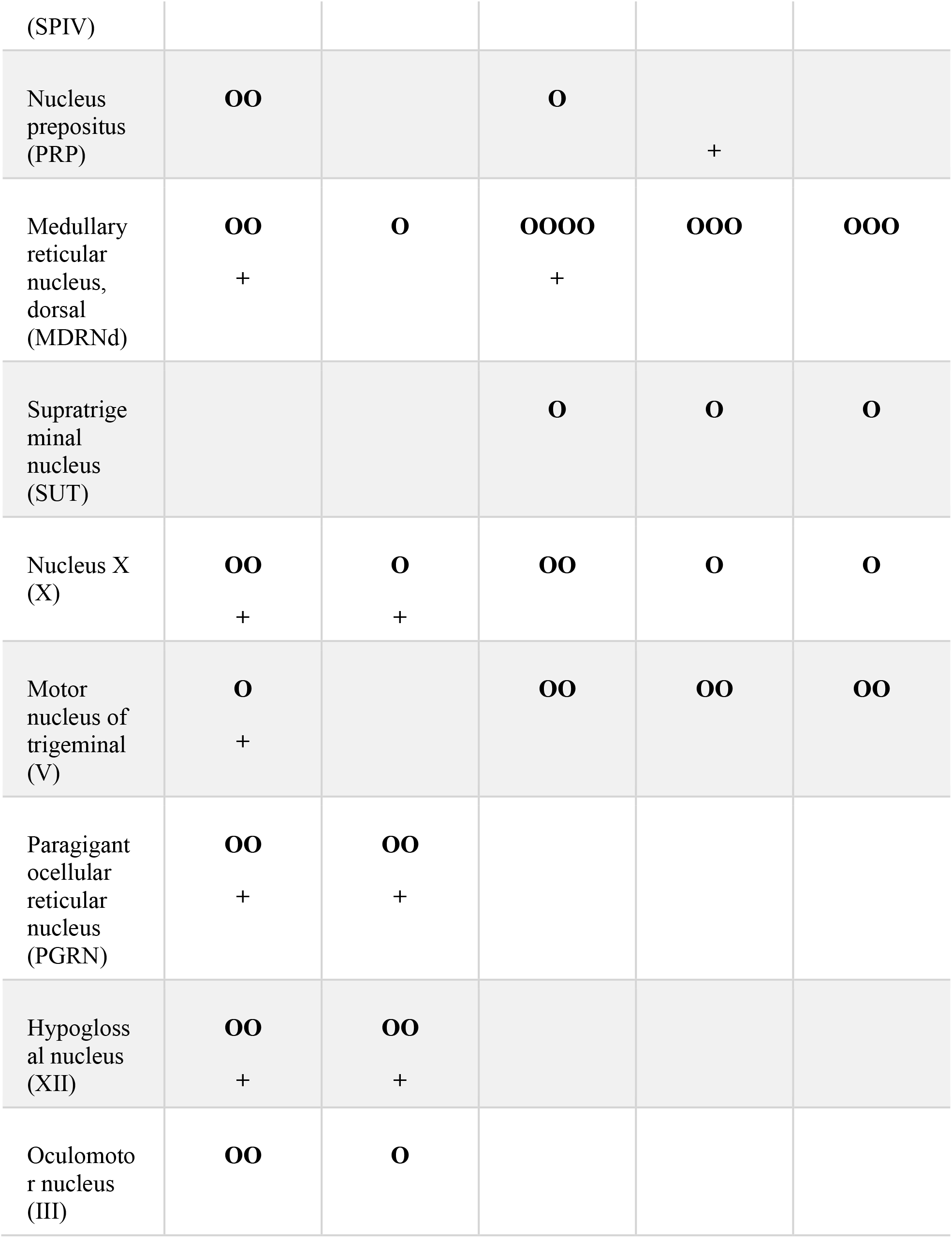
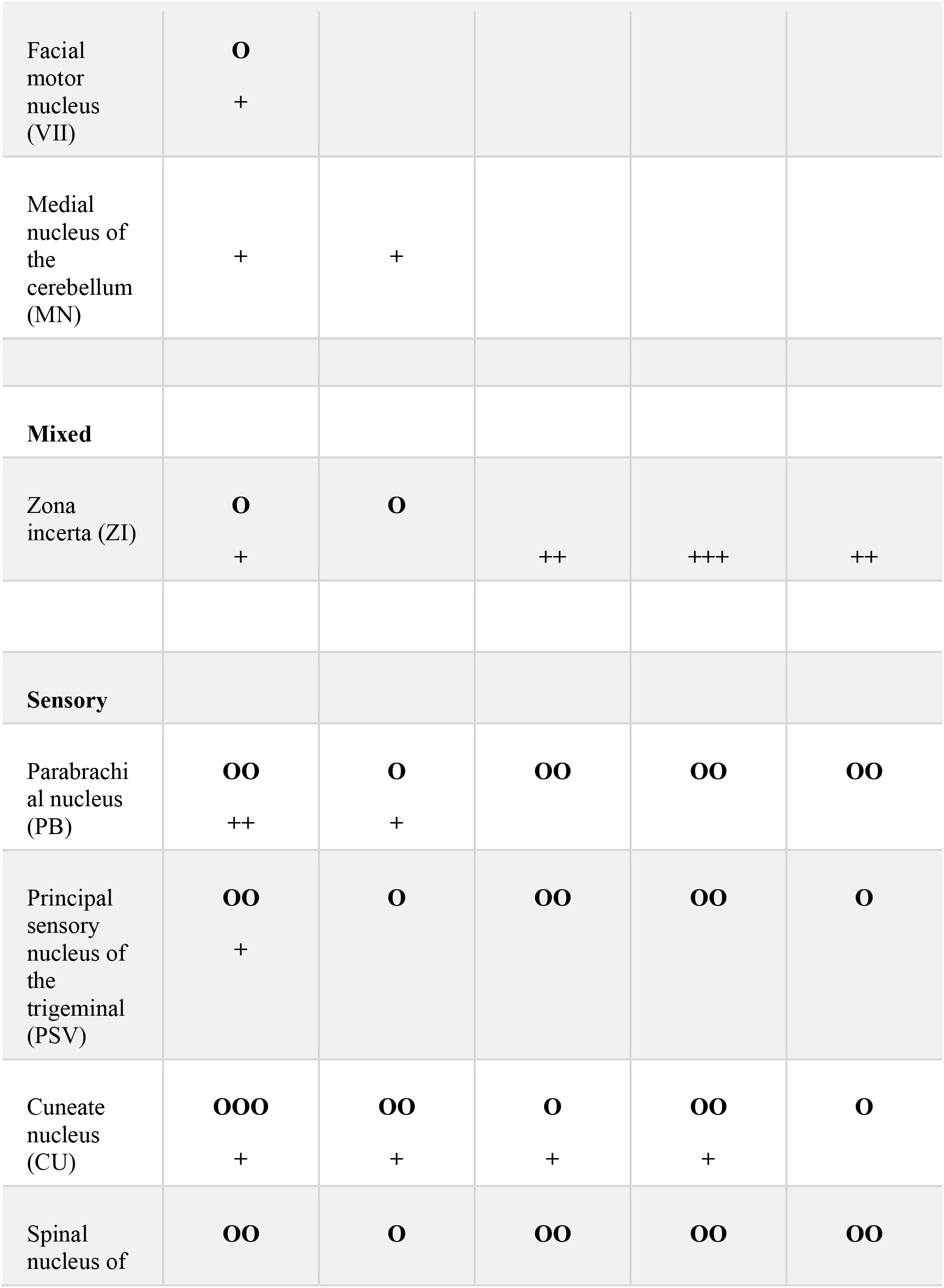
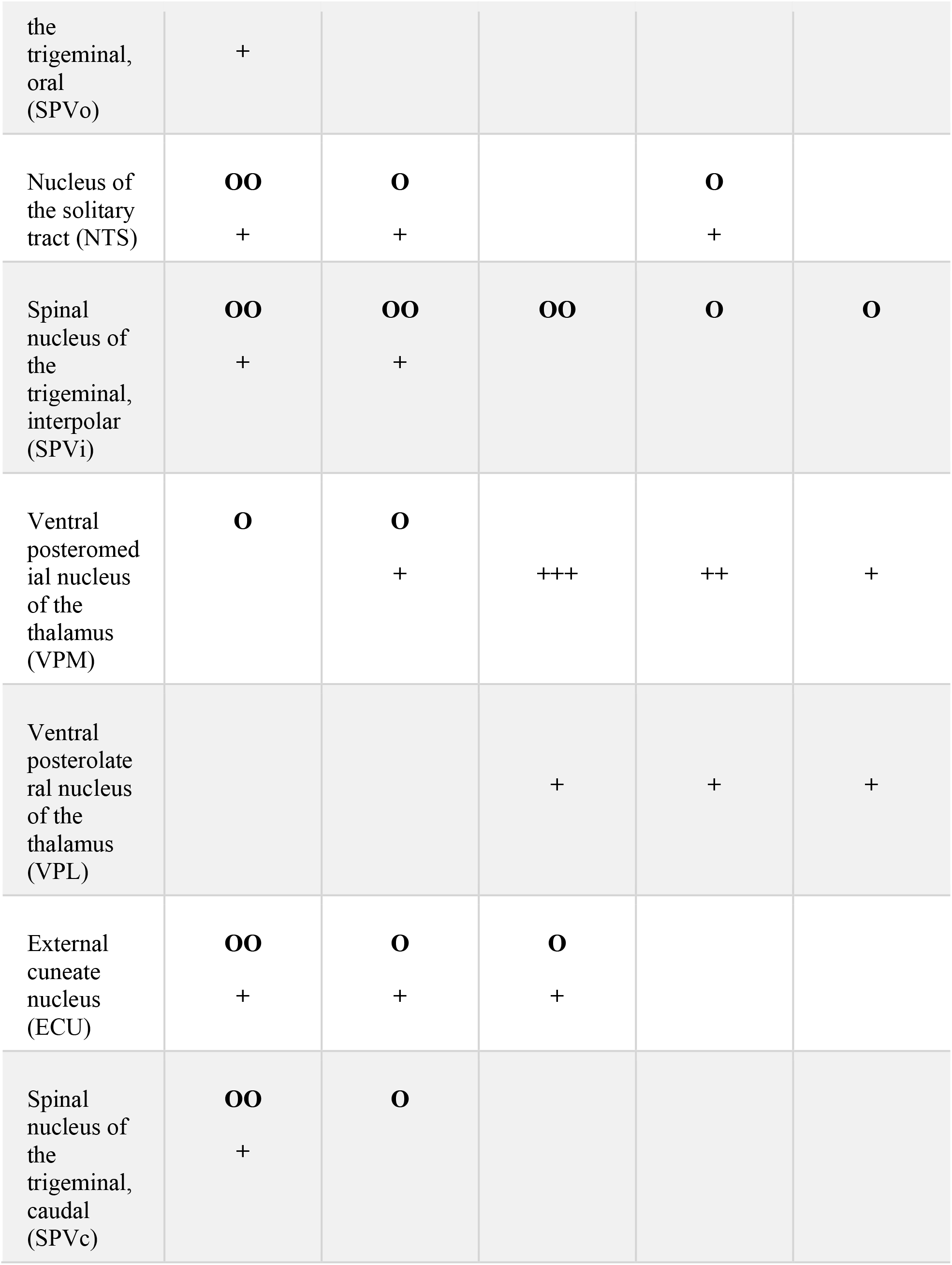
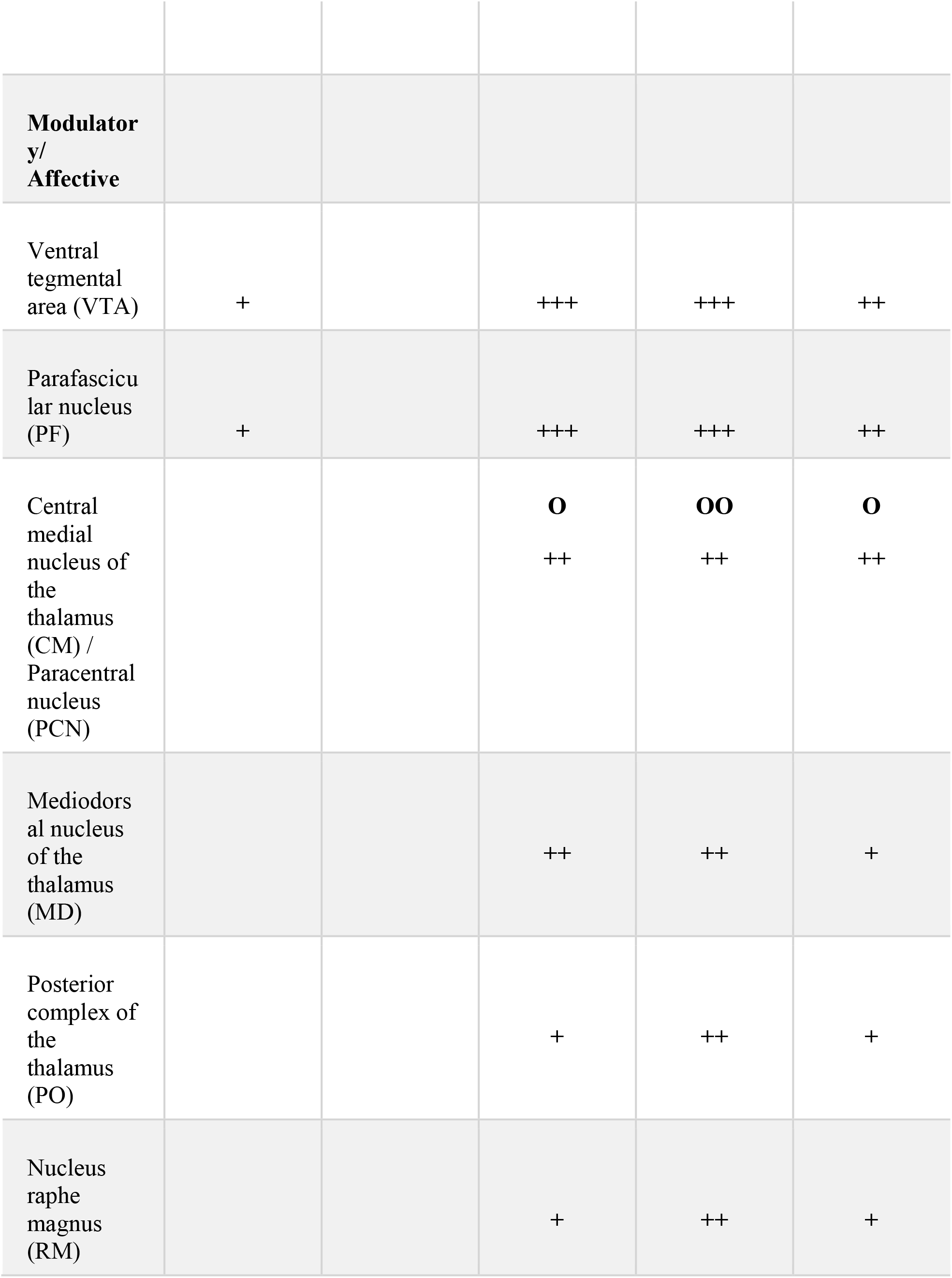
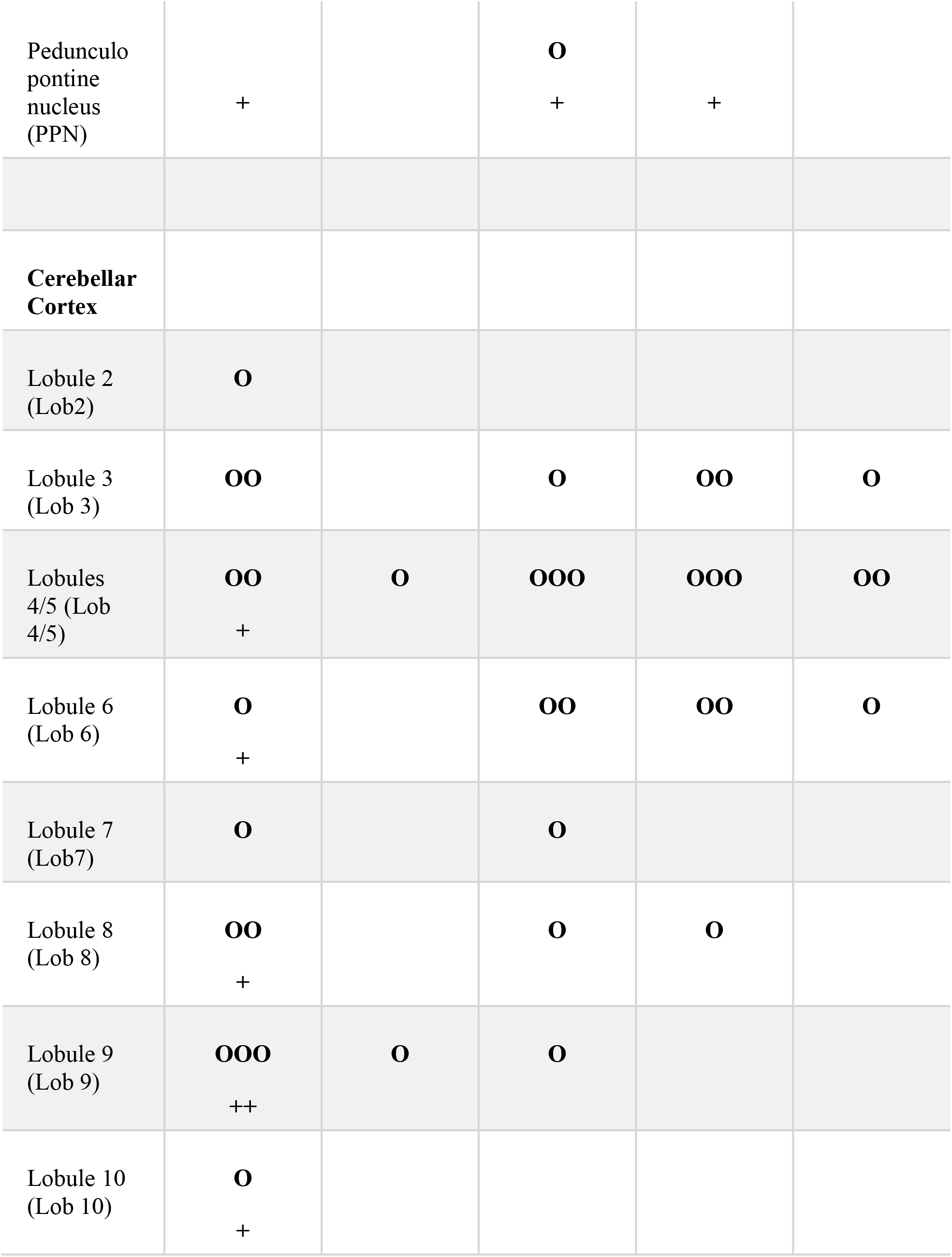
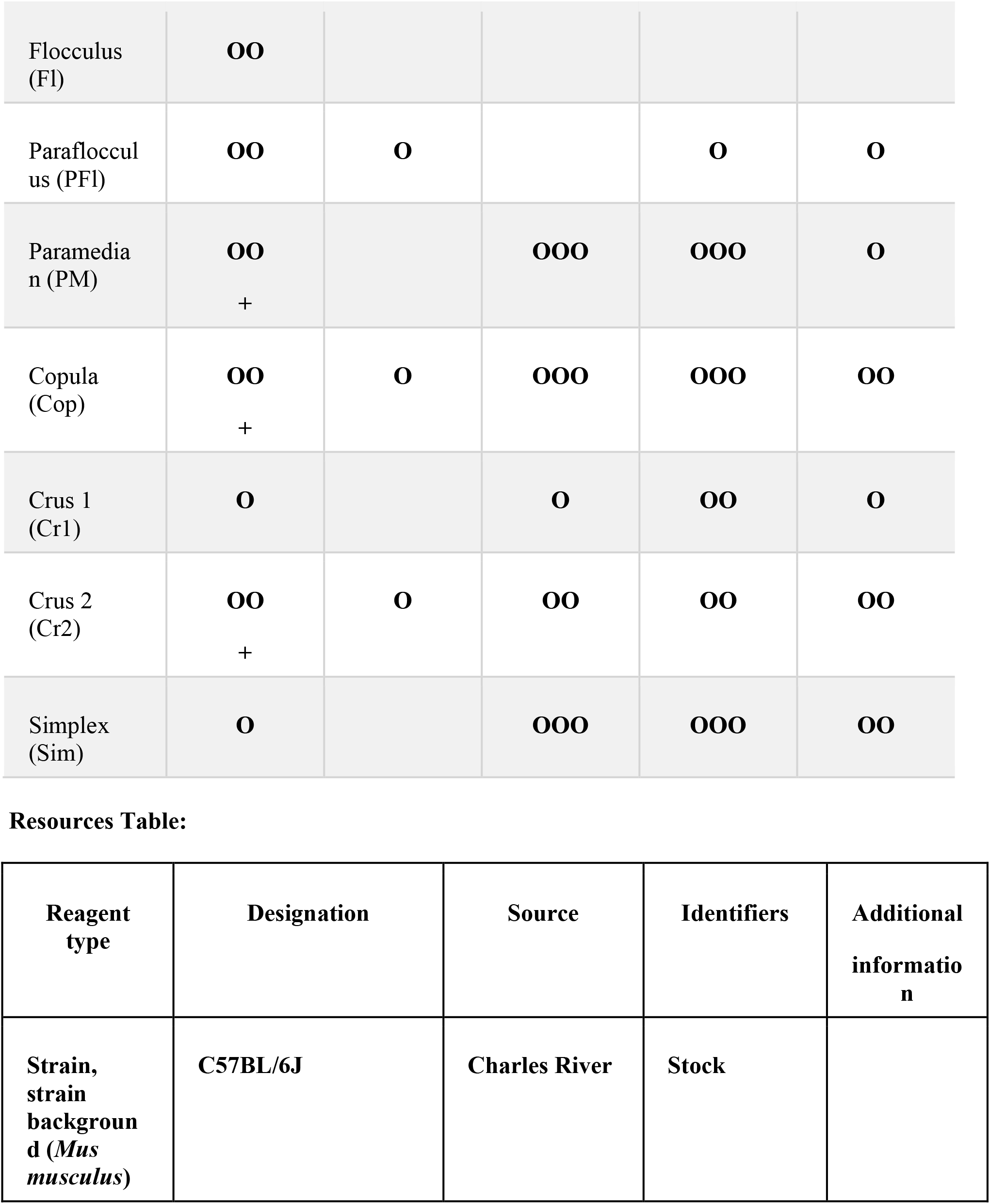
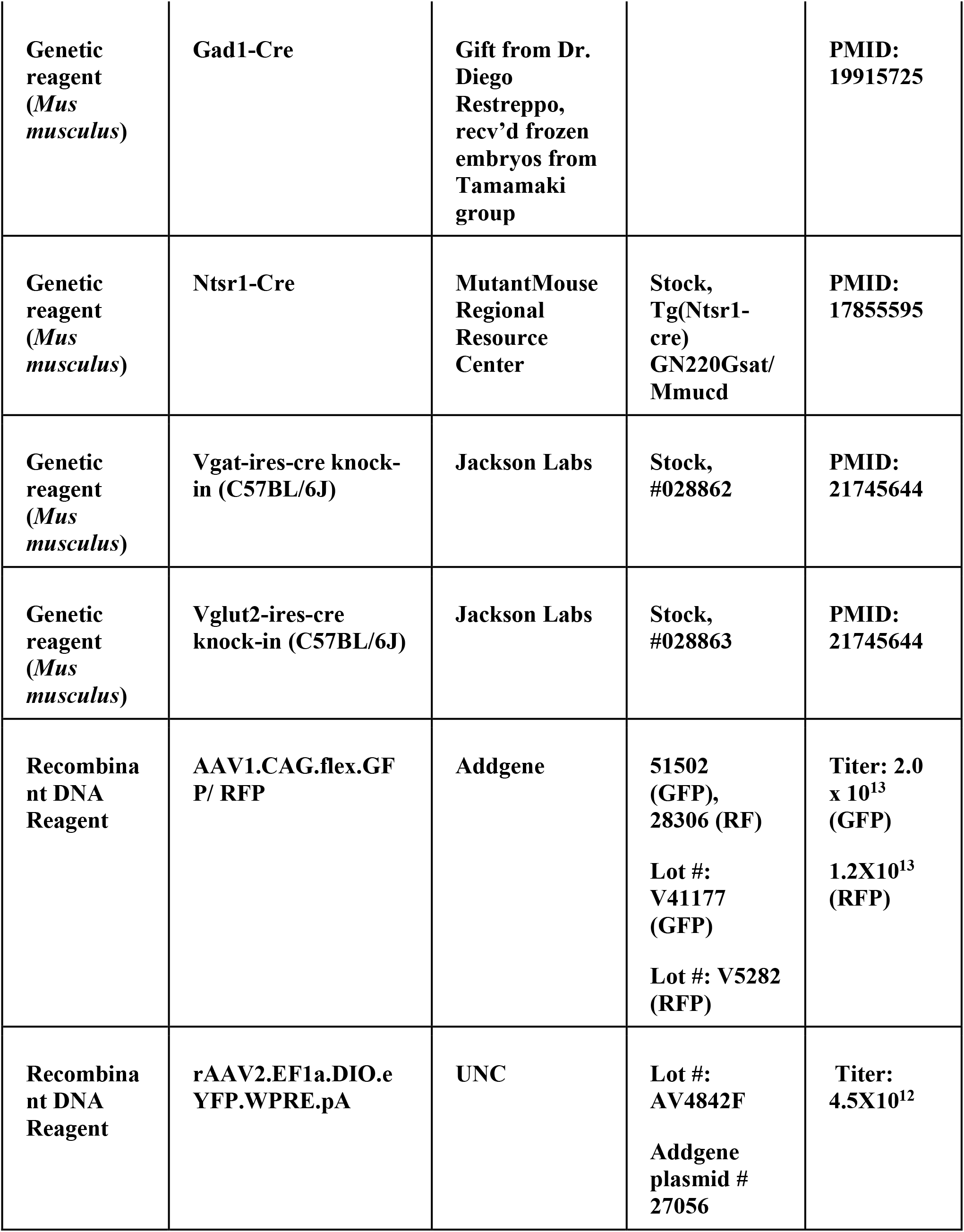
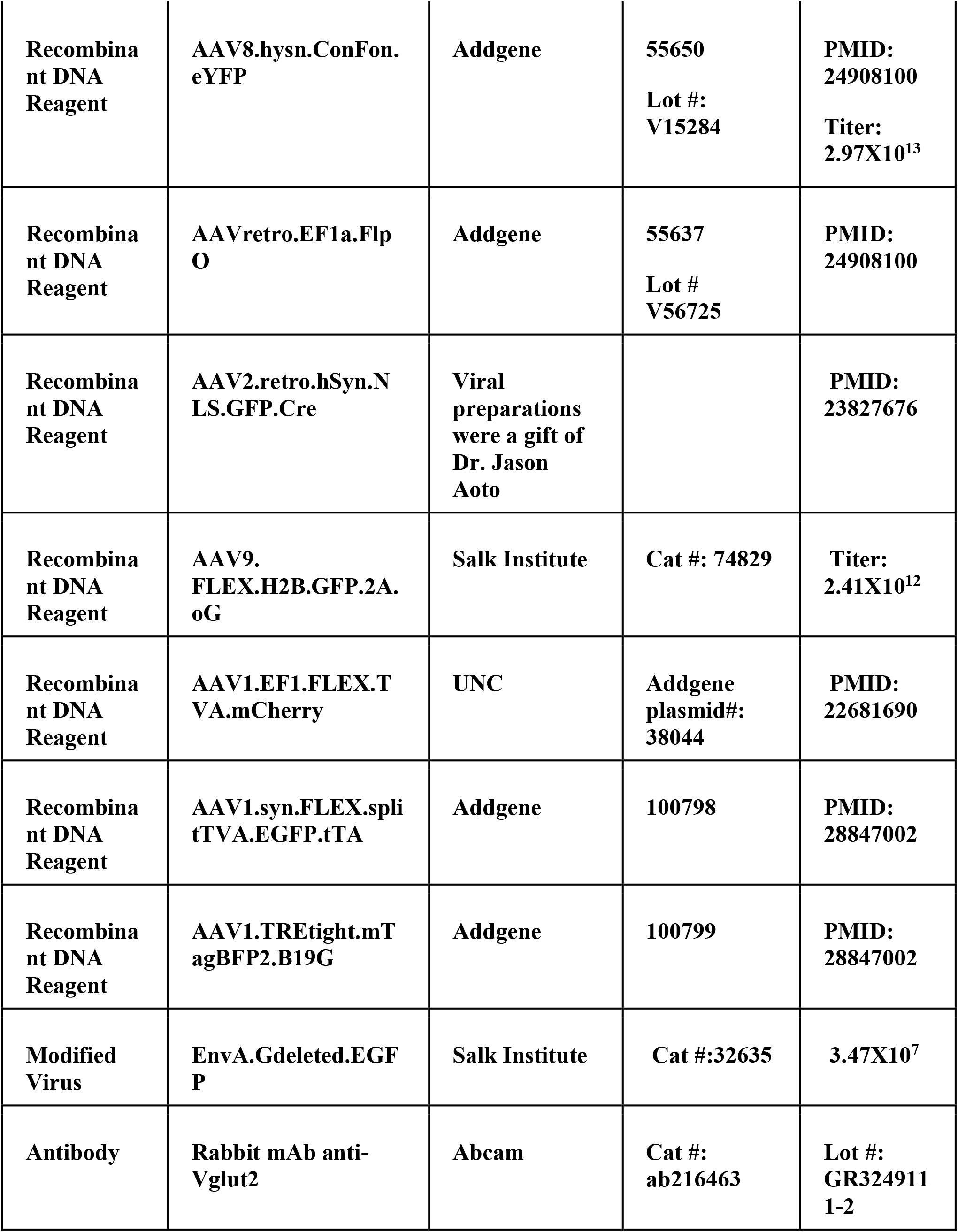
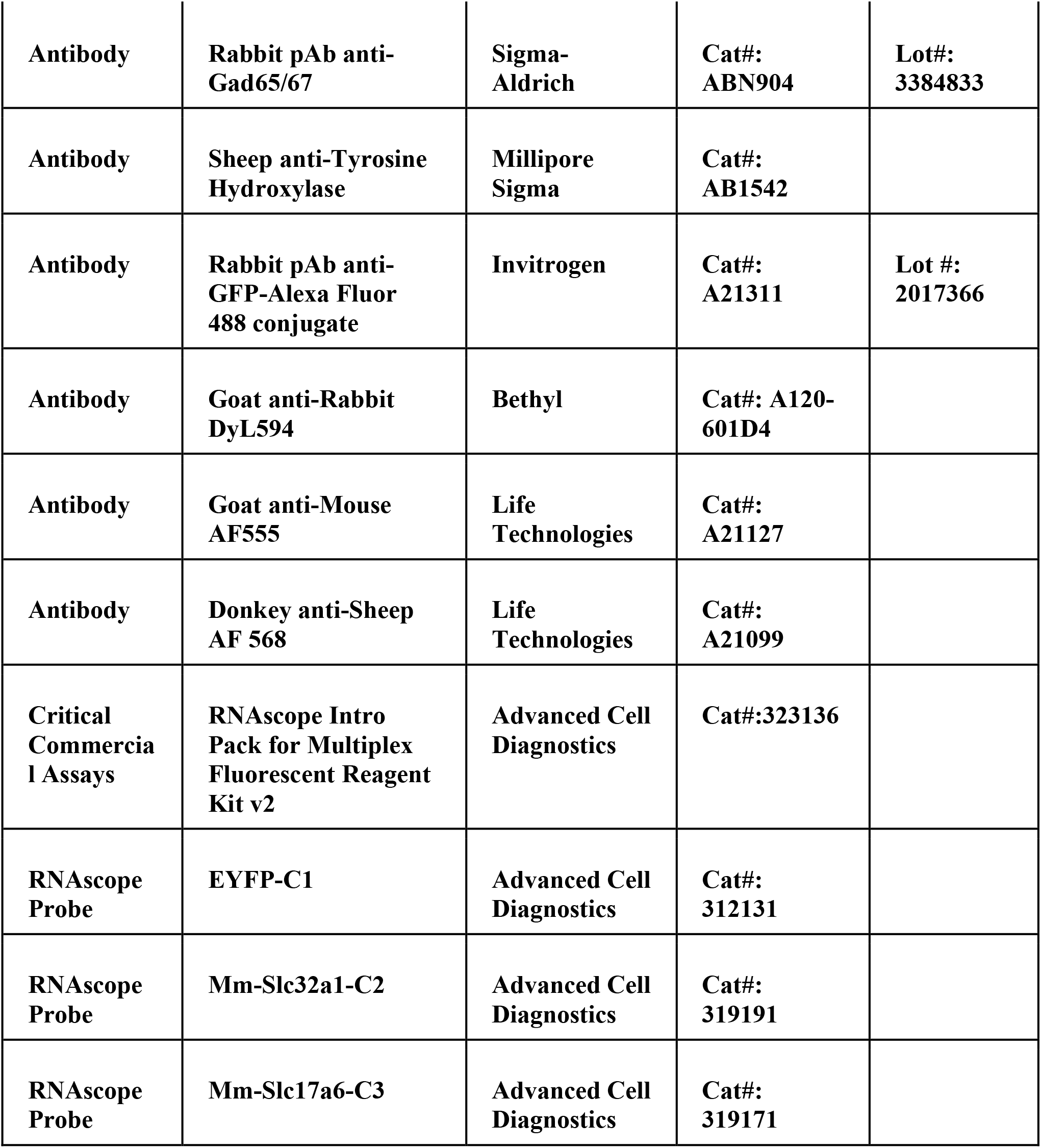
Anterograde tracing summary. Average RPS for all specimens in Int-Vgat (n = 6; including Allen specimen), Int^IO^-Vgat (n = 5), Int^RN^ (n = 4), Int-Ntsr1 (n = 6, including Allen specimen), and Int^RN^-Ntsr1 (n = 5). RPS depicted as symbols (+ for contralateral RPS, O for ipsilateral RPS). One symbol = avg RPS < 1, two symbols = avg RPS ≥ 1 and <2, three symbols = avg RPS ≥2 and <3, four symbols = avg RPS ≥ 3.

### Comparison of projection patterns across labeling methods

Across the distinct labeling methods, we observed a variety of notable patterns that differentiated them. First, Int cell sizes differed by Cre driver lines. We measured the cross-sectional area and elliptical diameter of somata of virally labeled cells. Int-Vgat neurons tended to be small with tortured dendrites (Fig. 5A-C; 14.4 ± 0.5 µm diameter, 109.3 ± 7.8 µm ^2^ area; N = 5 mice; n=316 neurons). By contrast, Int-Ntsr1 neurons were characteristically large with smooth dendrites (Fig. 5A-C; 22.2 ± 0.8 µm diameter; 224.7 ± 13.6 µm ^2^ area; N=5 mice; n=229 neurons). Similarly, Int^IO^-Vgat neurons were small (Fig. 5B-C; 14.5 ± 0.2 µm diameter; 103.4 ± 2.2 µm^2^ area, N = 5 mice; n=404 neurons;) and Int^RN^-Ntsr1 neurons larger (Fig. 5B-C; 22.1 ± 1.0 µm diameter; 238.2 ± 18.5 µm ^2^ area, N=4 mice; n=125 neurons). We compared these groups statistically and found the Int-Vgat and Int^IO^-Vgat cells were not significantly different from one another but were significantly smaller than Int-Ntsr1 and Int^RN^-Ntsr1 neurons (One-way ANOVA; Tukey’s multiple comparison test p<0.0001 for all cross-genotype comparisons of means across specimens; p>0.99 for all within genotype comparisons of means across specimens).

**Figure 5.**
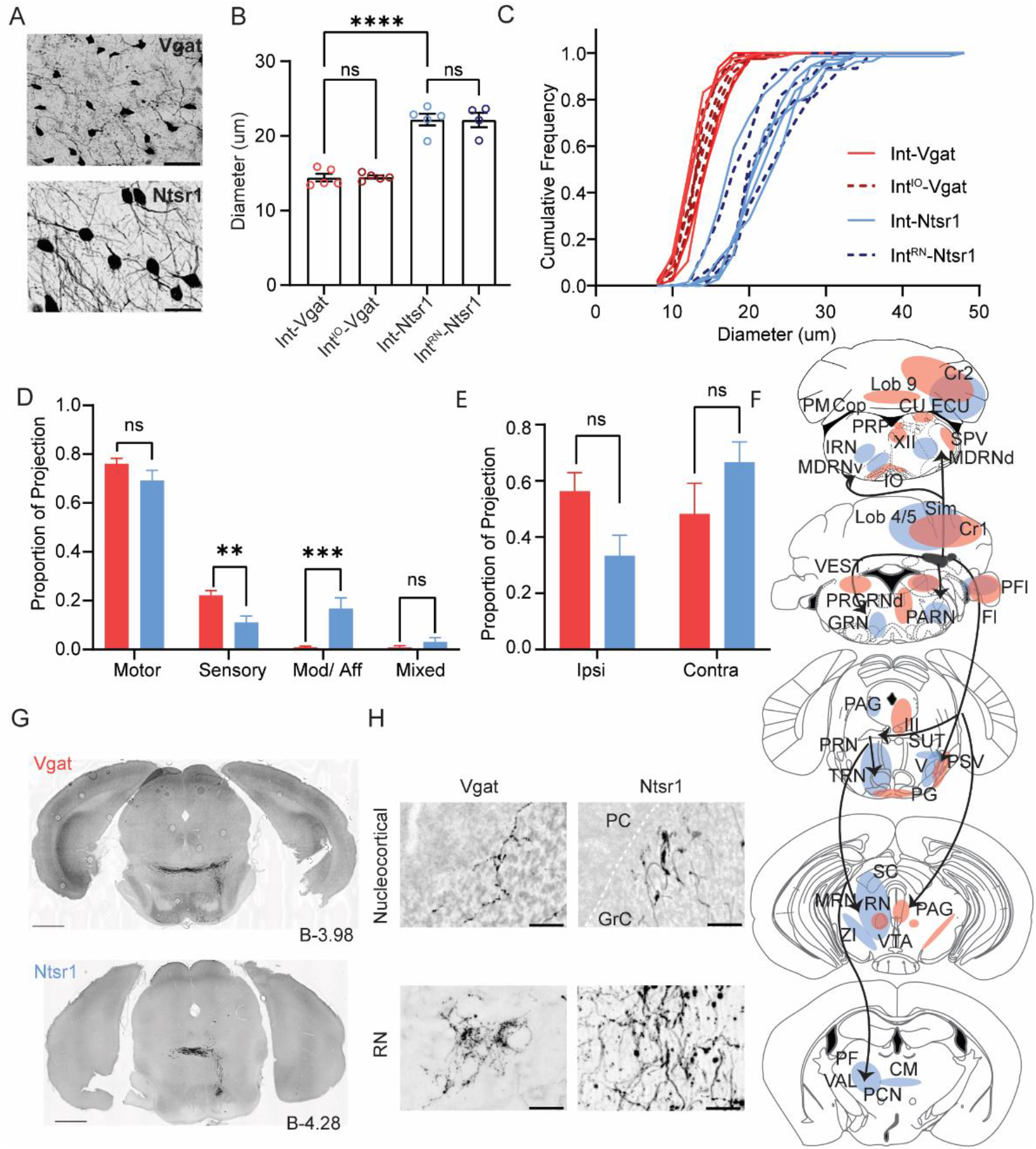
Comparison Int-Vgat and Int-Ntsr1 cell sizes and projection patterns. (A) Example YFP+ cells in a Vgat-Cre (top) and Ntsr1-Cre (bottom) specimen. Scale bars = 50 µms. (B) Differences in soma diameter of neurons based on isolation method. Grand mean ± SEM is plotted; per animal mean is denoted with colored circles (Int-Vgat =red, Int^IO^-Vgat = maroon, Ntsr1 = blue, Int^RN^-Ntsr1 = navy). Int-Vgat (n = 316 cells, 5 mice) or Int^IO^-Vgat neurons (n = 404 cells, 5 mice) are smaller than Int-Ntsr1(n = 229 cells, 5 mice) or Int^RN^-Ntsr1 neurons (n =125 cells, 4 mice; one-way ANOVA-Tukey’s multiple comparison’s test, P<0.0001, ****). (C) Cumulative frequency distribution of measured cell diameter for all specimens. (D) The average proportion of the total (summed) RPS value that is derived from projections to motor, sensory, or modulatory extracerebellar brain regions. Mean and SEM are plotted. Welch’s t-test with FDR correction of 1%, p = 0.035 (ns), 0.0014 (**), 0.00023 (***), 0.045 (ns) respectively. (E) Same as (D) but showing the contribution of ipsilateral or contralateral projections to total RPS per transgenic line. Welch’s t-test with FDR correction of 1%, p = 0.017 (ns) and 0.16 (ns), respectively. (F) Schematic of projection signatures from Ntsr1-Cre (blue) and Vgat-Cre (red). (G) Axons from Int-Vgat and Int-Ntsr1 follow unique paths through the pontine reticular nuclei (PRN). (H) Morphology differences in terminal contacts within the cerebellar cortex (top; boutons observed within the granule cell (GrC) layer; dotted white line in Nstr1 image denotes Purkinje Cell layer) and *red nucleus (RN*; bottom). Note mossy fiber nucleocortical terminals seen in Ntrs1-Cre mice but not Vgat-Cre mice. Scale bars = 50 µms. *Centromedial nucleus of the thalamus (CM), copula (Cop), Crus1 (Cr1) cuneate nucleus (CU), external cuneate nucleus (ECU), flocculus (Fl), gigantocellular reticular nucleus (GRN), hypoglossal nucleus (XII), inferior olive (IO), intermediate reticular nucleus (IRN), lateral reticular nucleus (LRN), lateral vestibular nucleus (LAV), medullary reticular nucleus, dorsal/ ventral subdivision (MDRNd/v), midbrain reticular nucleus (MRN), Nucleus Y (Y), nucleus prepositus (PRP), oculomotor nucleus (III), parabrachial (PB), paracentral nucleus of the thalamus (PCN) parafasicular nucleus of the thalamus (PF), paraflocculus (PFl), paragigantocellular reticular nucleus, dorsal (PGRNd), paramedian lobule (PM), parvicellular reticular nucleus (PARN), periaqueductal grey (PAG), pontine grey (PG), pontine reticular nucleus (PRN), principal sensory nucleus of the trigeminal (PSV), simplex lobule (Sim), spinal trigeminal nucleus, caudal/ interpolar subdivision (SPVc/i), spinal vestibular nucleus (SPIV), superior colliculus (SC), superior vestibular nucleus (SUV), supratrigeminal nucleus (SUT), superior vestibular nucleus (SUV), tegmental reticular nucleus (TRN), trigeminal motor nucleus (V), ventral tegmental area (VTA), ventromedial nucleus (VM), vestibular nuclei (VEST), zona incerta (ZI)*.

Second, we noted that many targets were distinct between genotypes and projection classes. We classified extracerebellar target regions as motor, sensory, and modulatory, based in part on groupings of the Allen Brain Atlas (see Methods). Notably, aggregate projection strength analyses indicated that on average, Int-Vgat neurons targeted sensory structures more densely than Int-Ntsr1 neurons (Fig. 5D; p = 0.001, unpaired Welch’s t-test). By contrast, we observed significantly stronger innervation of modulatory regions by Int-Ntsr1 than Int-Vgat (Fig. 5D; p= 0.0002, unpaired Welch’s t-test). Additionally, Int-Ntsr1 projections showed a contralateral bias and Int-Vgat an ipsilateral bias, but these trends were not significant when accounting for false positive discovery rates (Fig. 5E, F; p = 0.02, unpaired Welch’s t-test).

Third, qualitative assessment showed that axons tended to ramify in distinct subdivisions within the subset of targets shared by Int-Vgat and Int-Ntsr1. For example, Int-Vgat neurons projected to more lateral regions of the caudal spinal nucleus of the trigeminal (SPVc), and to more lateral and anterior divisions of the principle sensory nucleus of the trigeminal (PSV). Int-Ntsr1 projected to the medial edge of SPVc near the border with MDRNd/ PARN and to PSV near the border of the trigeminal (V). While both Int-Vgat and Int-Ntsr1 projected to the vestibular nuclei, Int-Vgat projections ramified more caudally in the spinal and medial nuclei than Int-Ntsr1. Int-Vgat projections to SC were absent. We also noted striking distinctions in the midbrain, where fibers from the two genotypes coursed in distinct locations. After decussating, Int-Vgat axons coursed farther lateral before turning ventrally toward the pontine nuclei. By contrast, Int-Ntsr1 axons turned ventrally at more medial levels after decussation, near the medial tracts through the pontine reticular nucleus (Fig. 5G). Because injection sites did not differ systematically across injection types, we interpret these distinctions to reflect targeting differences across cell classes.

Finally, as has been noted in previous studies, nucleocortical fiber morphology differs between excitatory and inhibitory neurons (Ankri et al., 2016; Houck and Person 2015; Gao et al., 2016; Batini et al., 1992). Int-Vgat injections labeled beaded varicosities devoid of mossy fiber morphological specializations, (Fig. 5H, top panels). Int-Ntsr1 labeled terminals with typical mossy fiber endings, large excrescences with fine filopodial extensions, and these predominantly targeted more intermediate lobules. Additionally, terminals in RN from Int-Vgat were very fine caliber while those from Int-Ntsr1 had thicker axons (Fig. 5H, bottom panels). While these observations are qualitative in nature, they align with the small cellular morphology of Int-Vgat neurons relative to Int-Ntsr1 neurons.

### Cell type specific input tracing using monosynaptic rabies virus

Having mapped pathways from diverse cell types of the intermediate cerebellar nuclei, we next investigated afferents to these cells (Fig. 6A). As described above, *in situ* hybridization in Vgat-Cre mice validated Vgat somatic expression in YFP labeled cells within the interposed nucleus, thus these mice were used for input tracing to inhibitory neurons (n = 3). However, although output tracing from Int-Ntsr1 was validated with immunolabel of terminals varicosities against Vglut2, *in situ* hybridization analysis of Ntsr1-Cre revealed 89% of YFP-positive cell bodies expressed Vglut2 probes (Fig. 3F, S3; 119/132 cells from 2 mice) but some labeled cells, possibly interneurons, expressed Vgat (15/132 cells in 2 mice). Thus, to ensure input mapping specific to excitatory neurons, we tested mRNA probe specificity of Vglut2-Cre mice (Fig. 6B, S3, Table S1): 178/179 YFP expressing cells (99%) expressed Vglut2 mRNA and 3/149 expressed Vgat probes. Therefore, Vglut2-Cre (n=3) mice were used to isolate inputs to glutamatergic Int populations. These mice were used in conjunction with modified rabies (EnvA-ΔG-Rabies-GFP/mCherry) and Cre-dependent receptor and transcomplementation helper viruses (Fig. 6A; see Methods; E. J. Kim et al., 2016; Wall et al., 2010; Watabe-Uchida et al., 2012; Wickersham et al., 2007, 2010). Direct rabies virus infection was limited to cells which expressed the receptor, TVA; transsynaptic jump was restricted by complementation of oG. In a subset of experiments, oG was restricted to TVA-expressing neurons (Liu et al., 2017). 72.9 ± 9.6% of starter cells in Vglut2-Cre specimens were mapped to Int (Fig. 6C,D); with the remaining starter cells located in the lateral (15%), medial (2%) and superior vestibular nucleus (5%). Similarly, 80.8 ± 4.7 % starter cells in Vgat-Cre mice were in Int, with the remainder in superior vestibular nuclei (7%), lateral (5%), medial (5%), and parabrachial (1%) nuclei (Fig. 6D, Table S3). Total numbers of starter cell estimates (Doykos et al., 2020), defined by presence of rabies and TVA (Fig. 6C) averaged 156 ± 131 in Vglut2-Cre and 307 ± 132 neurons in Vgat-Cre. TVA expression was not observed in cortex of Vgat-Cre or Vglut2-Cre mice, minimizing concerns of tracing contaminated by projections to cortical neurons.

**Figure 6.**
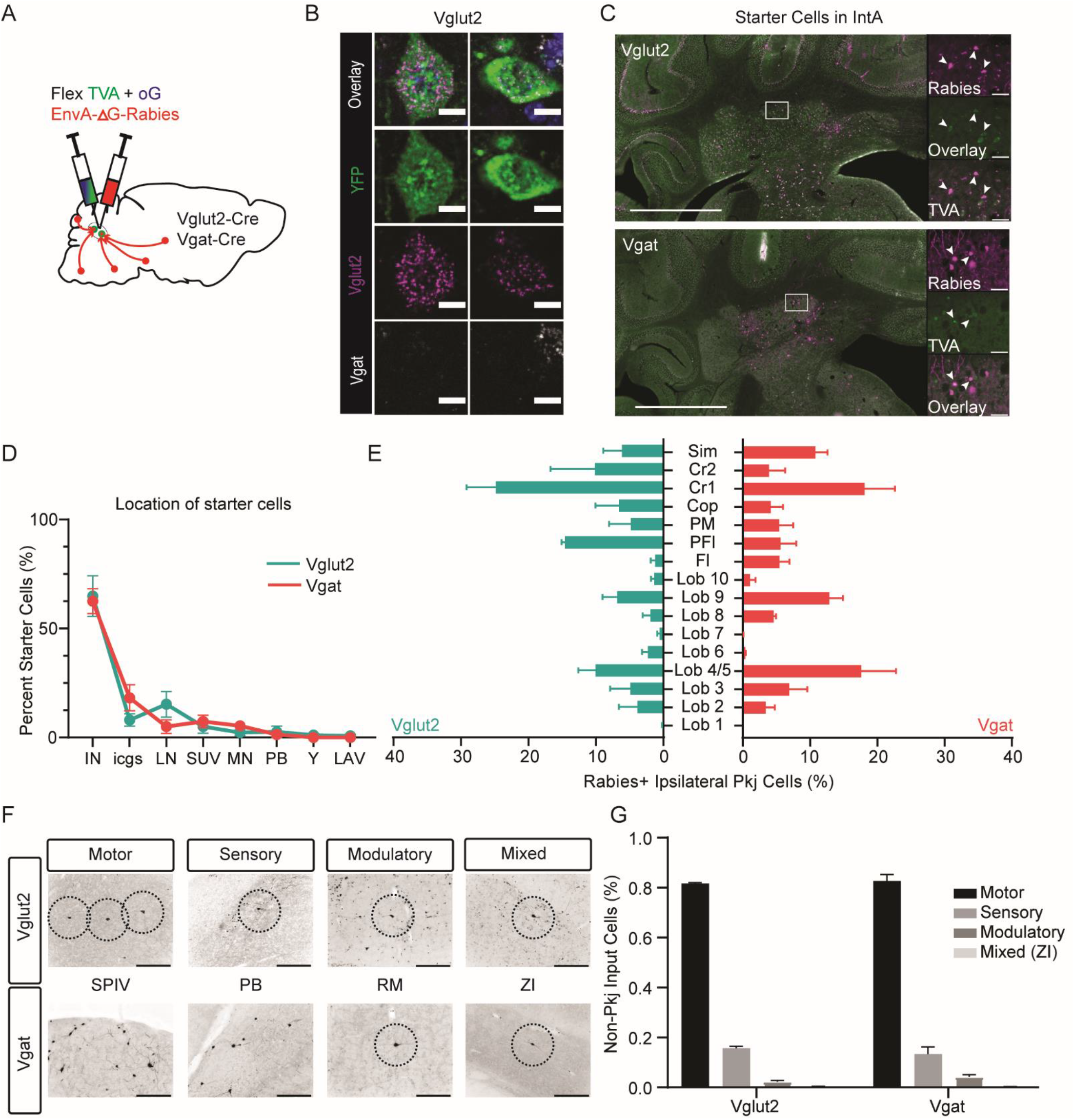
Monosynpatic tracing of inputs to Int-Vgat and Int-Vglut2. (A) Schematic of viral experiment. Cells labeled by this method provide monosynaptic input to Int. (B) Example Vglut2-YFP cells from *in situ* hybridization showing clear overlap with an mRNA probe against SLC17a6 (Vglut2) and no overlap with an mRNA probe against SLC32a1 (Vgat). Scale bars = 10 µms. (C) Example starter cells from both transgenic mouse lines in IntA. Scale bar = 1 mm. Insets to the right show Rabies (magenta, top), TVA (green channel, top), and overlay (bottom). Scale bar = 50 µms. (D) Locations of putative starter cells largely overlap for both cell types (mean + SEM). (E) Location of retrogradely labeled ipsilateral PCs by lobule. (F) Example extracerebellar rabies positive cells in motor (*spinal vestibular nuclei, SPIV*), sensory (*parabrachial, PB*), modulatory (*raphe magnus, RM*), and mixed (*zona incerta, ZI*) brain regions for both mouse lines. (G) Percent of non-PC inputs to Vglut2 or Vgat starter cells separated by modality. *Simplex lobule (sim), Crus1 (Cr1), Crus 2 (Cr2), Copula (cop), paramedian lobule (PM), Paraflocculus (PFl), Flocculus (FL)*.

The cerebellar nuclei receive a massive projection from Purkinje cells. The location of retrogradely labeled Purkinje cells (PC) was similar between specimen (Fig. 6E), regardless of genotype. PCs in ipsilateral Lobules 4/5, Crus 1, and Simplex were most densely labeled following rabies starting in both cell types. No contralateral PC label was observed in any specimen.

Extracerebellar input to Vglut2-Cre and Vgat-Cre cells was diverse and wide-ranging (Fig. S11). Both cell types receive input from brain regions related to motor, sensory, or modulatory functions (Fig. 6F), corroborating previous observations with traditional tracers (Fu et al., 2011; but see Barmack, 2003). For a complete list of brain regions which provide input to Vglut2-Cre and Vgat-Cre Int neurons, see Table S3 and Figure S9. Vglut2-Cre cells received a majority of inputs from ipsilateral sources, but not by large margins, with 64% of inputs originating in ipsilateral regions (36% contralateral). For Vgat-Cre cells, 54% of non-PC label was from ipsilateral sources (46% contralateral). These differences were not significant (p=0.2; unpaired Welch’s t-test). No extracerebellar region accounted for more than 10% of the total cells, suggesting widespread integration within Int. Of note, significantly more LRN neurons were retrogradely labeled following Vgat-Cre injections (5.9% of non-PC rabies labeled cells; >300 cells/specimen) than Vglut2-Cre (0.2% of total non-PC rabies label cells, <40/specimen p = 0.004 unpaired Welch’s t-test), suggesting a more extensive input to Int-Vgat neurons from LRN. Aside from this difference, extracerebellar projections to both cell types came from medial vestibular nuclei, TRN, and other reticular formation nuclei. We observed retrograde label in the contralateral medial cerebellar nucleus from both Vgat and Vglut2-Cre mice.

Many canonical sources of mossy fibers, such as ECU, PRN, TRN, PG, LRN (Parenti et al., 1996) were identified as sources of nuclear input as well as recipients of a projection from at least one cell type within Int (Fig. 7A-B; Tsukahara et al., 1983; Murakami et al., 1991). Figure 7B summarizes the inputs and outputs of both cell types ranked by average percentage of total rabies labeled cells within a given region (for retrograde tracing, excluding Purkinje cells) or average projection strength (for anterograde tracing), excluding the weakest projections. The only brain regions which received a projection but were not also retrogradely labeled were the thalamic nuclei. In converse, the only brain regions with retrogradely labeled cells but not anterograde projections were motor cortex, somatosensory cortex, subthalamic nucleus, and lateral hypothalamus, among other minor inputs (Table S3).

**Figure 7.**
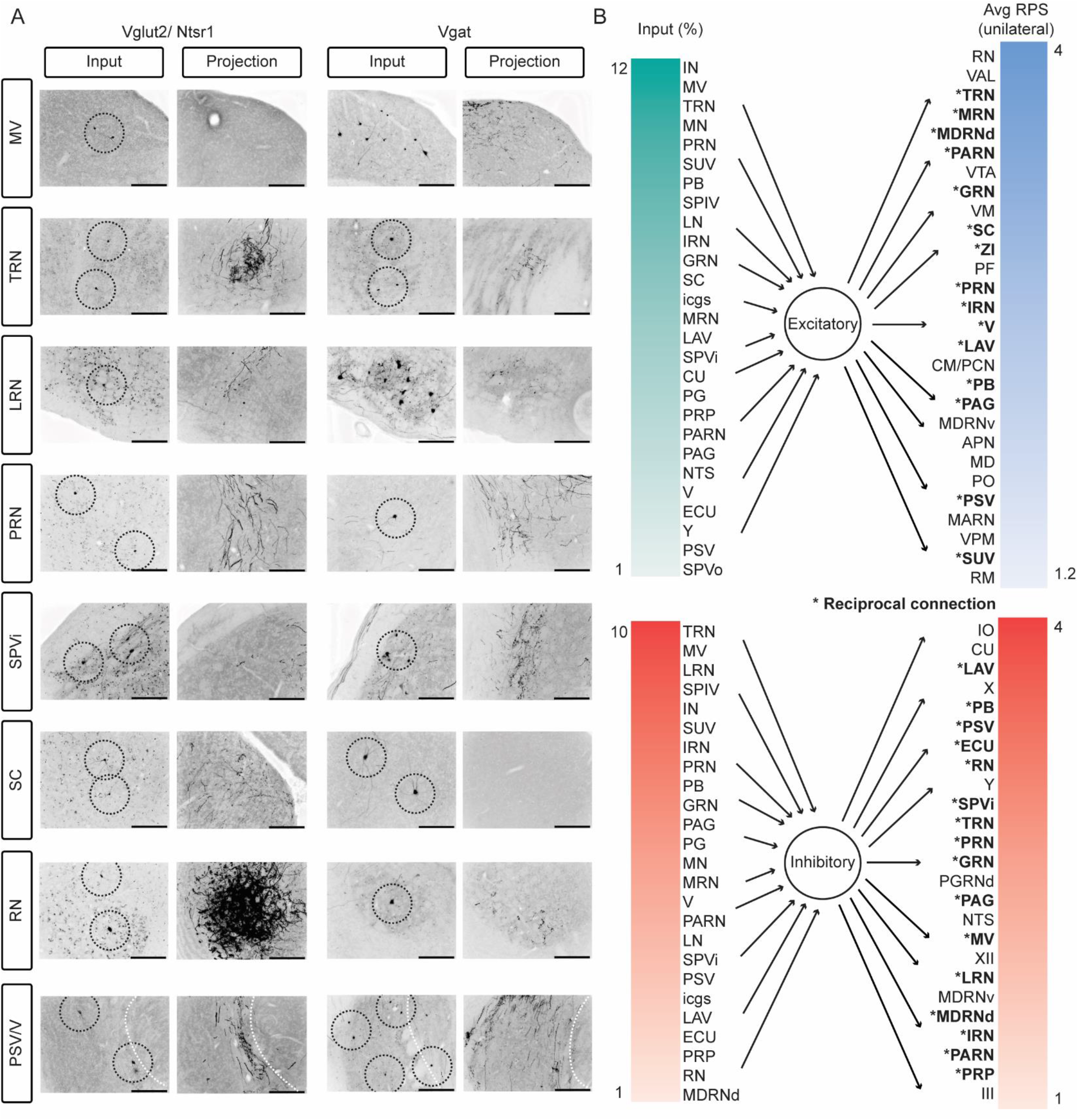
Reciprocal loops between Int and extracerebellar targets, for both Vglut2 and Vgat expressing cells. (A) Images depicting rabies labeled cells (columns 1 and 3, rabies + cells circled if singular or very small) and projections that included axon varicosities to the same regions at the same coordinates relative to bregma (columns 2 and 4). *Medial vestibular nuclei (MV), tegmental reticular nucleus (TRN), lateral reticular nucleus (LRN), pontine reticular nuclei (PRN), spinal trigeminal nucleus, interpolar subdivision (SPVi), superior colliculus (SC), red nucleus (RN), principal sensory nucleus of the trigeminal (PSV), motor nucleus of the trigeminal (V).* White dotted line denotes boundary between PSV and V. (B) Inputs and outputs listed in order of percent of non-PC rabies labeled cells (left) and relative projection strength (right). Only inputs with greater than 1% of total extracerebellar rabies labeled cells and regions with mean relative projection strengths greater than 1 are listed. Asterisks denote regions that constituted a major afferent (>1% total input) and received a major projection (an RPS >1 in Vgat-Cre mice and >1.2 in Ntsr1-Cre mice). *Anterior pretectal nucleus (APN), centromedial nucleus of the thalamus (CM), cuneate nucleus (CU), external cuneate nucleus (ECU), gigantocellular reticular nucleus (GRN), hypoglossal nucleus (XII), inferior olive (IO), intermediate reticular nucleus (IRN), interposed nucleus (IN), interstitial cell groups (icgs), lateral nucleus (LN), lateral reticular nucleus (LRN), lateral vestibular nucleus (LAV), magnocellular reticular nucleus (MARN), medial nucleus (MN), medial vestibular nuclei (MV), mediodorsal nucleus of the thalamus (MD), medullary reticular nucleus, dorsal/ ventral subdivision (MDRNd/v), midbrain reticular nucleus (MRN), motor nucleus of the trigeminal (V), nucleus of the solitary tract (NTS), nucleus raphe magnus (RM), nucleus X (X), Nucleus Y (Y), nucleus prepositus (PRP), oculomotor nucleus (III), parabrachial (PB), paracentral nucleus of the thalamus (PCN), parafasicular nucleus of the thalamus (PF), paragigantocellular reticular nucleus, dorsal (PGRNd), paramedian lobule (PM), parvicellular reticular nucleus (PARN), periaqueductal grey (PAG), pontine grey (PG), pontine reticular nucleus (PRN), principal sensory nucleus of the trigeminal (PSV), red nucleus (RN), spinal trigeminal nucleus, interpolar/ oral subdivision (SPVi/o), spinal vestibular nucleus (SPIV), superior colliculus (SC), superior vestibular nucleus (SUV), supratrigeminal nucleus (SUT), superior vestibular nucleus (SUV), tegmental reticular nucleus (TRN), trigeminal motor nucleus (V), ventral anterior-lateral complex of the thalamus (VAL), ventral posteromedial nucleus of the thalamus (VPM), ventral tegmental area (VTA), ventromedial nucleus (VM), zona incerta (ZI)*.

## Discussion

Here we systematically examined the input and output patterns of diverse cell populations of the interposed cerebellar nucleus, Int, using intersectional viral tracing techniques. Consistent with previous work, we found that the putative excitatory output neurons of Int collateralize to regions of the contralateral brainstem, spinal cord and thalamus and more restrictedly to the caudal ipsilateral brainstem, including to regions recently shown to control forelimb musculature. However, we also found that Int GABAergic projection neurons innervate brainstem regions other than IO, including the pontine nuclei, medullary reticular nuclei, and sensory brainstem structures. Interestingly, IO-projecting neurons collateralize to comprise, in part, these projections. Inputs to these distinct cell types were also mapped using monosynaptic rabies tracing. We found that inputs to glutamatergic and Vgat neurons of the intermediate cerebellum are largely similar with only the lateral reticular nucleus standing out as preferentially targeting Vgat neurons. Merging anterograde and retrograde datasets, region-level reciprocal loops between Int and brainstem targets were similar across both cell types.

The most surprising results were the diverse projections of GABAergic neurons of Int. To address concerns that these projections may be the result of a methodological artifact, we note a variety of data that support our interpretation. First, *in situ* hybridization and immunolabel support the view that Vgat-Cre is restricted to Vgat expressing neurons that express Gad65/67 in terminal boutons. Second, projection patterns of excitatory neurons were distinct, particularly within the ipsilateral caudal brainstem and diencephalon, thus non-specific viral label cannot account for the data. Third, AAV-retroCre injections into RN – a putative target of both Int-Vgat and Int-Ntsr – labeled targets matching mixed projections of excitatory and inhibitory neurons, including terminal label in IO. Finally, we used an intersectional approach, targeting Vgat-Cre expressing neurons that project to the IO. This method of isolating Int inhibitory neurons also consistently labeled terminals elsewhere in the brainstem. Taken together, leak of Cre cannot explain the sum of these observations.

Another study restricting tracer to lateral (dentate) nucleus Sox14-Cre expressing neurons, a transcription factor marking nucleo-olivary neurons, showed terminal label in the IO as well as the oculomotor nucleus (III). Based on retrograde tracing from III, terminals there were interpreted to reflect virus uptake by nucleus Y near the injection site (Prekop et al., 2018). This finding raised the question of whether brainstem and midbrain targets of Int-Vgat neurons described in the present study are merely a consequence of viral uptake in regions neighboring the interposed nucleus. Although projections were more extensive following larger injections in Vgat-Cre mice, we observed axon varicosities outside IO following injections that were completely restricted to the interposed nucleus. As has been noted in previous studies, the ventral border of the interposed nucleus is poorly distinguished but houses numerous islets of cells within the white matter tracts (Sugihara and Shinoda, 2007; Sugihara 2011). These regions receive Purkinje input from zebrin negative zones, and have been proposed to be distinct subregions of the cerebellar nuclei. A medial population, named the interstitial cell group, resides ventrally between the medial and interposed nuclei. An anterior extension, the anterior interstitial cell group, resides ventral to the interposed nucleus, and more posterior and laterally, the parvocellular interposed and lateral cell groups, neighboring nucleus Y, complete this constellation of loosely organized cell groups. Our data hint that these regions may house Vgat neurons that produce more extensive extra-IO projections, including nucleocortical beaded axons distinct from nucleocortical mossy fibers, although this conjecture will remain speculative until methodological advances permit cell type specific tracing from such minute regions to be carried out.

While these inhibitory projections were unknown, these data, combined with previous literature from the medial nucleus, suggest that inhibitory projections from the cerebellar nuclei may be a more prominent circuit motif than is widely appreciated. The medial nucleus contains glycinergic projection neurons that innervate ipsilateral brainstem nuclei matching contralateral targets of excitatory neurons (Bagnall et al., 2009). Additional evidence of inhibitory outputs includes dual retrograde tracing suggesting that nucleo-olivary projections from medial nucleus and the vestibular complex collateralize to the ventromedial hypothalamic nucleus (Diagne et al., 2001; Li et al., 2017). Studies combining retrograde horse radish peroxidase tracing from the basilar pontine nuclei (i.e. pontine gray) with immunohistochemistry observed double labeled GABA immunopositive neurons in the lateral nucleus of rats and cats (Aas & Brodal, 1989), although the literature is inconsistent (Schwarz & Schmitz, 1997). More recent work in mice tracing Vgat-Cre neurons of the lateral nucleus listed projections to a variety of brainstem structures as well as IO (Locke et al., 2018), but these results were not discussed.

Despite these corroborating experimental results, we note that our data may appear to contradict conclusions drawn from a dual-retrograde tracing study, in which only minor dual retrograde label was observed in the lateral and interposed nuclei following tracer injections into IO and RN or IO and TRN (Ruigrok & Teune, 2014). This study concluded that two distinct populations exist within the CbN: one which projects widely to several regions and one which projects exclusively to IO. However, this study did report a small number of cells colabeled by retrograde injections to IO and TRN as well as IO and RN. This observation may account for the present finding that a population of neurons which projects to both IO and premotor nuclei exists in smaller numbers, and that topographic specificity may have precluded previous methods from fully detecting the collateralization of inhibitory populations.

Projection patterns of glycinergic medial and vestibular nucleus neurons have an ipsilateral bias relative to excitatory contralateral projections. (Bagnall et al., 2009; Prekop et al., 2018; Sekirnjak et al., 2003; Shin et al., 2011). This organizational structure has been proposed to potentially mediate axial muscular opponency. While there was a trend for an ipsilateral targeting of Int-Vgat neurons, this bias was not significant when accounting for false discovery rates (p=0.02), with both excitatory and inhibitory cells projecting bilaterally. Future studies investigating the functional roles of these projections may explore agonist/antagonist opponency in motor targets of these projections, which remain lateralized for limb musculature. Additionally, the widespread observation of Purkinje neurons that increase rates during cerebellar dependent behaviors may suggest the potential for a double disinhibitory pathway through the cerebellar nuclei, if these Purkinje neurons converged on inhibitory nuclear output neurons (De Zeeuw and Berrebi, 1995; De Zeeuw, 2020).

What might be the role of inhibitory projections from the cerebellar nuclei? Two intriguing patterns emerged that are suggestive of potential function. First, inhibitory projections targeted more sensory brainstem structures than excitatory outputs. Predicting sensory consequences of self-generated movement, termed forward models, is a leading hypothesis for the role of cerebellum in sensorimotor behaviors. While populations of Purkinje neurons may perform this computation, it is unknown how forward models are used by downstream targets. Inhibitory projections from cerebellum to sensory areas would seem to be ideally situated to modulate sensory gain of predicted sensory consequences of movement (Brooks et al., 2015; Shadmehr, 2020). Moreover, negative sensory prediction error could be used to actively cancel predicted sensory reafference (Kim et al., 2020; Requarth & Sawtell, 2014; Shadmehr, 2020; Conner et al., 2021), raising implications for a combined role of negative sensory prediction error in guiding learning both through modulation of climbing fiber signaling in IO and through modulation of sensory signals reaching the cerebellum upon which associative learning is built. Second, GABAergic projections to the pontine nuclei, which are themselves a major source of mossy fiber inputs to the cerebellum, suggests a regulatory feedback pathway that could operate as a homeostat akin to the feedback loops through the IO (Medina et al., 2002). The pontine nuclei are a major relay of cortical information into the cerebellum. Thus, through inhibitory feedback, cortical information could potentially be gated to facilitate strategic (i.e. cortical) control for novel skill learning, or turned down to facilitate automatic (i.e. ascending/non-cortical) control of movements (Schwarz and Thier, 1999).

The present study compliments a recent collection of papers examining cerebellar nuclear cell types. Transcriptomics analyses of the cerebellar nuclei identified three distinct excitatory cell types within IntA. These classes included two broad projection types: those that target a wide array of brainstem nuclei and those that target the ZI (Kebschull et al., 2020). Another recent study identified two distinct interposed cell types based on projection patterns to the spinal cord, which were shown to constitute a minority of neurons (<20%). Nevertheless, these spinal-projecting neurons collateralized to many other targets, including the MDRNv, RN, and the VAL (Sathyamurthy et al., 2020). Inhibitory projections were not examined in these studies, thus it will be interesting to examine how the inhibitory projection neurons identified in the present study map onto transcript clusters of the inhibitory cell types, 5 total across the cerebellar nuclei. At a minimum, these clusters would include IO-projecting neurons, interneurons, MN glycinergic projection neurons, and a collateralizing population of inhibitory neurons identified here (Ankri et al., 2015; Bagnall et al., 2009; Fujita et al., 2020; Zoé Husson et al., 2014; Kebschull et al., 2020; Sathyamurthy et al., 2020).

Inputs to these neuronal populations were largely similar, though we observed minor differences in the input signatures of Int-Vglut2 and Int-Vgat. Many more neurons in the lateral reticular nucleus (LRN) were labeled following Vgat-Cre starting cells for monosynaptic rabies tracing, suggesting a predominant innervation of inhibitory neurons by LRN. It remains unclear if there are differences in input connectivity between Vgat+ subgroups, specifically interneurons and projection neurons, or whether Gad65/67 expressing neurons co-express GlyT2. In comparing input and outputs to diverse cell types, we noticed that reciprocal loops were common, broadening themes of reciprocal loops demonstrated previously (Tsukahara et al., 1983, Beitzel et al., 2017), to also include inhibitory neurons. Such loops resemble neural integrators used in gaze maintenance or postural limb stabilization (Albert et al., 2020; Cannon & Robinson, 1987), another potential functional role of the anatomy presented here. Interestingly, we observed neocortical inputs to the intermediate groups of the cerebellar nuclei. We speculate that these regions may conform to the reciprocal loop motif, albeit polysynaptically, predicting that thalamic targets innervate neocortical areas that project back to the cerebellar nuclei.

In conclusion, the anatomical observations presented here open the door to many potential functional studies that could explore the roles of inhibitory projections in real-time motor control, sensory prediction and cancellation, and dynamic cerebellar gain control, as well as explore roles of afferents to the cerebellar nuclei such as the motor cortex. Taken together, the present results suggest distinct computational modules within the interposed cerebellar nuclei based on cell types and shared, but likely distinct, participation in motor execution.

## Materials and Methods

### Animals

All procedures followed the National Institutes of Health Guidelines and were approved by the Institutional Animal Care and Use Committee at the University of Colorado Anschutz Medical Campus. Animals were housed in an environmentally controlled room, kept on a 12 h light/dark cycle and had ad libitum access to food and water. Adult mice of either sex were used in all experiments. Genotypes used were C57BL/6 (Charles River Laboratories), Neurotensin receptor1-Cre [Ntsr1-Cre; MutantMouse Regional Resource Center, STOCK Tg(Ntsr1-cre) GN220Gsat/ Mmucd], Gad1-Cre (Higo et al., 2009); Vgat-Cre[#028862]; Jackson Labs], Vglut2-Cre [#028863; Jackson Labs]. All transgenic animals were bred on a C57BL/6 background. Gad1 and Ntsr1-Cre mice were maintained as heterozygotes and were genotyped for Cre (Transnetyx). For all surgical procedures, mice were anesthetized with intraperitoneal injections of a ketamine hydrochloride (100 mg/kg) and xylazine (10 mg/kg) cocktail, placed in a stereotaxic apparatus, and prepared for surgery with a scalp incision.

### Viral injections

Injections were administered using a pulled glass pipette. Unilateral pressure injections of 70-200 nl of Cre-dependent reporter viruses (AAV1.CAG.flex.GFP; AAV2.DIO.EF1a.eYFP; AAV8.hysn-ConFon.eYFP, see Key Resources Table) were made into Int. Injections were centered on IntA, with minor but unavoidable somatic label appearing in posterior interposed (IntP), lateral nucleus (LN), interstitial cell groups (icgs) and the dorsal region of the vestibular (VEST) nuclei, including dorsal portions of the superior vestibular nucleus (SUV), lateral vestibular nucleus (LAV), and Nucleus Y (Y). We occasionally observed minor somatic label in the parabrachial nucleus (PB) and the cerebellar cortex (Cb-Ctx) anterior or dorsal, respectively, to Int in Vgat injections. In control injections (n = 3; virus in C57/Bl6 mice or off-target injection into Ntsr-1 Cre mice), viral expression was not detected. We did not see appreciable somatic label in the medial nucleus (MN) of any specimens. For RN injections, craniotomies were made unilaterally above RN (from bregma: 3.5 mm, 0.5 mm lateral, 3.6 mm ventral).

For Int injections, unilateral injections were made at lambda: 1.9 mm posterior, 1.6 mm lateral, 2.1 mm ventral. For IO injections, the mouse’s head was clamped facing downward, an incision was made near the occipital ridge, muscle and other tissue was removed just under the occipital ridge, and unilateral injections were made at 0.2 mm lateral, and 2.1 mm ventral with the pipet tilted 10° from the Obex. This method consistently labeled IO and had the advantage of avoiding accidental cerebellar label via pipette leakage. To achieve restricted injection sites, smaller volumes were required in Vgat-Cre mice compared to Ntsr1-Cre mice (40-100 nL vs 150-200 nL, respectively). The smallest Vgat-Cre injection was made iontophoretically using 2 M NaCl. Current (5 μA) was applied for 10 mins, the current was removed and after a waiting period of 5 mins, the pipet was retracted. Retrograde labeling of RN-projecting IntA neurons was achieved through AAVretro-EF1a-cre (Tervo et al., 2016). Retrograde injections of RN were performed simultaneously with flex-GFP injections of IntA. Retrograde virus (AAVretro-EF1a-Flp) was injected to IO one week before reporter viruses because of the different targeting scheme and mice were allowed to heal one week prior to the reporter virus injection. All mice injected with AAVs were housed postoperatively for ∼ 6 weeks before perfusion to allow for viral expression throughout the entirety of the axonal arbor. Control injections were performed where Cre or Flp expression was omitted, either by performing the injections in wild type mice or in transgenic mice without the Retro-flp injection into IO or RN, confirming the necessity of recombinase presence in reporter expression (Fenno et al., 2017).

For monosynaptic rabies retrograde tracing, AAV1-syn-FLEX-splitTVA-EGFP-tTA and AAV1-TREtight-mTagBFP2-B19G (Addgene; Liu et al., 2017) were diluted 1:200 and 1:20, respectively, and mixed 1:1 before co-injecting (100 nL of each; vortexed together) unilaterally into IntA of Vgat-IRES-Cre (n = 3) and Vglut2-IRES-Cre (n =1) mice. Two additional Vglut2-IRES-Cre mice were prepared using AAV1.EF1a.Flex.TVA.mCherry (University of North Carolina Vector Core; (Watabe-Uchida et al., 2012)) and AAV9.Flex.H2B.GFP.2A.oG (Salk Gene Transfer, Targeting and Therapeutics Core; (E. J. Kim et al., 2016)). After a 4-6-week incubation period, a second injection of EnvA.SADΔG.eGFP virus (150-200 nL) was made at the same location (Salk Gene Transfer, Targeting and Therapeutics Core; (E. J. Kim et al., 2016; Wall et al., 2010; Wickersham et al., 2007). Mice were sacrificed one week following the rabies injection and prepared for histological examination. Control mice (C57Bl/6; n = 1) were injected in the same manner, however, without Cre, very little putative Rabies expression was driven, though 8 cells were noted near the injection site. No cells were identified outside this region (Supplementary Figure 6).

### Tissue Preparation and imaging

Mice were overdosed with an intraperitoneal injection of a sodium pentobarbital solution, Fatal Plus (MWI), and perfused transcardially with 0.9% saline followed by 4% paraformaldehyde in 0.1 M phosphate buffer. Brains were removed and postfixed for at least 24 hours then cryoprotected in 30% sucrose for at least 24 hours. Tissue was sliced in 40 μm consecutive coronal sections using a freezing microtome and stored in 0.1 M phosphate buffer. Sections used for immunohistochemistry were floated in PBS, permeabilized using 0.1-0.3% Triton X-100, placed in blocking solution (2-10% Normal Goat serum depending on antibody) for 1-2 hours, washed in PBS, and incubated in primary antibodies GFP (1:400), Gad65/67 (1:200), Vglut2(1:250), and TH (1:200) for 24-72 hours. Sections were then washed in PBS thrice for a total of 30 mins before incubation in secondary antibodies (Goat anti Rabbit DyL594, Goat anti-Mouse AF555 (1:400), or Goat anti-Sheep AF568 (1:400), see Key Resources Table) for 60-90 mins. Finally, immunostained tissue was washed in PBS and mounted in Fluoromount G (SouthernBiotech).

Every section for rabies experiments and every third section for anterograde tracing experiments was mounted onto slides and imaged. Spinal cord sections were also sliced in 40 μm consecutive coronal sections with every 4^th^ section mounted. Slides were imaged at 10x using a Keyence BZX-800 microscope or a slide-scanning microscope (Leica DM6000B Epifluorescence & Brightfield Slide Scanner; Leica HC PL APO 10x Objective with a 0.4 numerical aperture; Objective Imaging Surveyor, V7.0.0.9 MT). Images were converted to TIFFs (OIViewer Application V9.0.2.0) and analyzed or adjusted via pixel intensity histograms in Image J. We inverted fluorescence images using greyscale lookup tables in order to illustrate results more clearly. YFP terminals stained for neurotransmitter transport proteins (Gad65/67 or Vglut2) were imaged using a 100X oil objective on a Marianas Inverted Spinning Disc confocal microscope (3I). We imaged in a single focal plane of 0.2 μm depth to analyze colocalization of single terminal endings. Images were analyzed in Image J.

### Analysis of overlap by genetically defined neurons

To distinguish overlap of Cre expression with transmitter markers, we performed *in situ* hybridization. For *in situ* hybridizations (ISH), RNAse free PBS was used for perfusions and the tissue was cryoprotected by serial applications of 10, 20, and 30% Sucrose for 24-48 hours each. The brain tissue was then embedded in OCT medium and sliced to 14 μm thick sections on a cryostat (Leica HM 505 E). Tissue sections were collected directly onto SuperFrostPlus slides and stored at −80 deg for up to 3 months until RNA *in situ* hybridization for EYFP (virally driven), SLC32a1 (Vgat), and SLC17a6 (Vglut2) from RNAscope Multiplex Fluorescent Reagent Kit v2 (Advanced Cell Diagnostics). The slides were defrosted, washed in PBS and baked for 45 mins at 60°C in the HybEZ™ oven (Advanced Cell Diagnostics) prior to post-fixation in 4% PFA for 15 mins at 4°C and dehydration in ethanol. The sections were then incubated at room temperature in hydrogen peroxide for 10 mins before performing target retrieval in boiling 1X target retrieval buffer (Advanced Cell Diagnostics) for 5 mins. The slides were dried overnight before pretreating in protease III (Advanced Cell Diagnostics) at 40°C for 30 mins. The RNAscope probes #312131, # 319191, #319171 were applied and incubated at 40°C for two hours. Sections were then treated with preamplifier and amplifier probes by applying AMP1, AMP2 at 40°C for 30 min and AMP3 at 40°C for 15 min. The HRP signals were developed using Opal dyes (Akoya Biosciences): 520 (EYFP probe), 570 (Vglut2 probe), and 690 (Vgat probe) and blocked with HRP blocker for 30 mins each.

The cerebellar nuclei were stained using DAPI for 30 seconds before mounting in Prolong Gold (ThermoFisher). Washes were performed twice between steps using 1X wash buffer (Advanced Cell Diagnostics). Fluorescence was imaged for YFP, Vglut2, Vgat, and DAPI using a Zeiss LSM780 microscope. Each image was captured using a 34-Channel GaAsP QUASAR Detection Unit (Zeiss) at 40X magnification in water from 14 µm sections. Images were stitched using ZEN2011 software and analyzed in ImageJ. Cre-expressing cells were identified by somatic labeling in the YFP channel; colocalization with the Vgat or Vglut2 channel was determined by eye using a single composite image and the “channels tool”. Positive (Advanced Cell Diagnostics, PN 320881) and negative control probes (Advanced Cell Diagnostics, PN 32087) resulted in the expected fluorescent patterns (Fig. S3).

We analyzed the fidelity of our transgenic lines using virally mediated YFP somatic label and DAPI staining to identify cells expressing Cre and analyzed the colocalization of Vgat or Vglut2 mRNA within the bounds of a YFP cell. The YFP signal was often less punctate than our other endogenous mRNA probes, thus we restricted our analysis to cells largely filled by YFP signal that contained DAPI stained nuclei. Due to the high expression patterns of Vgat and Vglut2, analyzing by eye was reasonable. Only a total of 4 cells across all analyzed sections appeared to have ISH-dots in both the Vgat and Vglut2 channels. This may be due to poor focus on these individual cells or background fluorescence. Two sections per animal and one (Vglut2-IRES-Cre and Gad1-Cre) or two animals (Ntsr1-Cre and Vgat-IRES-Cre) per transgenic line were counted.

In preliminary studies, we tested a Gad1-Cre driver line (Higo et al., 2009) for specificity since Gad1 was recently identified as a marker of inhibitory neurons within the cerebellar nuclei (Kebschull et al., 2020). However, in this line, we observed clear instances of both Gad65/67 and Vglut2-immunoreactivity in YFP labeled terminal varicosities as well as some Vglut2 mRNA expression in YFP expressing somata (Table S1, Fig. S3-4). 87% YFP expressing cells colocalized with Vgat (60/69 cells) and 13% colocalized with Vglut2 (9/60 cells). The clear instances of promiscuity in Gad-1 Cre mice precluded further use of these mice in the present study.

### Cell size analysis

We imaged cells within IntA at 20x then used the “Measure” tool in ImageJ to gather the cross-sectional area and the “Fit ellipse” measurement to gather minimum and maximum diameter which we converted from pixels to microns using reference scale bars. We report the maximum diameter. We analyzed 15-110 well focused and isolated cells for each specimen. Statistical analyses were conducted using a repeated-measures analysis of variance (ANOVA) on the per-animal and grand means of cell diameter per experimental condition.

### Brain region classification

We used a combination of the Allen Mouse Brain Reference Atlas and the *Mouse Brain in Stereotaxic Coordinates* by Franklin and Paxinos to identify brain regions, while noting that there were minor differences in location, shape and naming of the brain regions between these reference sources (Lein et al., 2007; Franklin & Paxinos, 2008). We generally grouped the dorsolateral and anterior subdivisions of Int because they were often co-labeled, are difficult to confidently distinguish, and occur at similar anterior-posterior (AP) coordinates. For monosynaptic rabies tracing and difficulty in targeting multiple viruses to the exact same location-we grouped all subdivisions of the interposed nucleus (IN; anterior, posterior, dorsolateral). In general, we followed nomenclature and coordinates respective to bregma of the Allen Mouse Brain Reference Atlas including its classification conventions of motor, sensory, modulatory/ affective sources from the 2008 version. Thalamic regions were classified as motor if they project to motor cortices; sensory if they project to sensory cortices, with intralaminar thalamic nuclei classified as modulatory/ affective. The intermediate and deep layers of the superior colliculus harbored terminal fields and retrogradely labeled neurons and is thus classified as motor.

### Projection quantification

Following viral incubation periods, we mapped terminals to a collection of extracerebellar targets spanning the anterior-posterior (A-P) axis from the posterior medulla to the thalamus. We assigned terminal fields a semi-quantitative relative projection strength (RPS) of 0-4 based on the density and anterior-posterior spread (Table 1). The values were assigned relative to the highest density projection target for each genotype: All Ntsr1-Cre projection fields were assessed relative to the density of terminals in RN whereas Vgat-Cre specimens were assessed relative to the density of IO terminals (Fig. S1). Briefly, a terminal field that was both dense and broad (in spanning the anterior-posterior axis) was assigned a relative projection strength (RPS) of 4, semi-dense and semi-broad assigned a 3, semi-dense and/ or semi-broad a 2, and fields determined to be neither dense nor broad but nevertheless present, were assigned an RPS of 1. In addition, we compared our specimens to analogous preparations published in the Allen Mouse Brain Connectivity Atlas, specifically the histological profile of Cre-dependent labeling following injections into IntA of either Ntsr1-Cre or Slc32a1(Vgat)-ires-Cre mice. These publicly available sources recapitulated projection signatures from lab specimens (Table S1). We included the Allen injection data in our analysis of average projection strength for Ntsr1-Cre (n=1) and Vgat-Cre (n=1) specimen but did not use the histological images of these injections here. The full histological profiles of genetically restricted GFP label from the Allen can be found at: 2011 Allen Institute for Brain Science. Mouse Brain Connectivity Atlas. Available from: http://connectivity.brain-map.org/, experiments #264096952, #304537794.

We determined the average proportion of the total RPS value that is derived from specific projections (to specific modalities or hemispheres; Fig. 5D, E) by summating the RPS values to every region receiving a projection per specimen. We then divided this number by summated RPS values in the groupings of interest. We report the average proportion of total RPS values across all specimens in each experimental cohort. These measurements are therefore indicative of the strength of projection to certain modalities or hemispheres, and not simply a measure of the number of brain regions targeted.

### Rabies quantification

We identified presumptive starter cells as rabies (mCherry) positive cells that also contained GFP (AAV1-syn-FLEX-splitTVA-EGFP-tTA). We used an antibody against EGFP (see Resources Table) to visualize TVA at these concentrations, but mBFP (AAV1-TREtight-mTagBFP2-B19G) could not be visualized. However, G expression is restricted to cells expressing TVA due to the necessity of the tetracycline transactivator gene encoded by the virus delivering TVA (Liu et al., 2017). In two Vglut2 rabies mice, we identified presumptive starter cells as rabies positive cells within the cerebellar nuclei where both mCherry (AAV1.EF1.Flex.TVA.mCherry) and GFP (AAV9.Flex.H2B.GFP.2A.oG.GFP/ EnvA.SADΔG.eGFP) were expressed. We could not easily identify cells in which all three components were present due to overlapping fluorescence from the oG and modified rabies viruses, thus starter cell identification is an estimate (Doykos et al., 2020). An additional caveat in these two Vglut2 rabies mice is that we are unable to distinguish starter cells from local interneurons infected transsynaptically which may artificially inflate the number of starter cells in these specimens.

## Abbreviations

APN: Anterior Pretectal Nucleus
CbCtx: Cerebellar Cortex
CbN: Cerebellar Nuclei
CM: Centromedial nucleus of the thalamus
CN: Cochlear Nucleus
CU: Cuneate Nucleus
CUN: Cuneiform Nucleus
DTN: Dorsal Tegmental Nucleus
ECU: External Cuneate Nucleus
GoC: Golgi Cells
GRN: Gigantocellular Reticular Nucleus
IC: Inferior Colliculus
III: Oculomotor Nucleus
IN: Interposed Nucleus
IntA: Anterior Interposed Nucleus
IO: Inferior Olive
IRN: Intermediate reticular nucleus
LAV: Lateral vestibular Nucleus
LC: Locus Coeruleus
LDT: Lateral dorsal tegmental nucleus
LN: Lateral Cerebellar Nucleus
LRN: Lateral Reticular Nucleus
MARN: Magnocellular reticular nucleus
MD: Mediodorsal nucleus of the thalamus
MDRNd: Medullary reticular nucleus dorsal
MDRNv: Medullary reticular nucleus ventral
MLI: Molecular Layer Interneurons
MN: Medial Cerebellar Nucleus
MRN: Midbrain reticular nucleus
MV: Medial vestibular nucleus
NLL: nucleus of the lateral lemniscus
NTS: Nucleus of the solitary tract
PAG: Periaqueductal grey
PARN (PCRt): Parvicellular reticular nucleus
PAS: Parasolitary nucleus
PB: Parabrachial nuclei
PC: Purkinje Cells
PCG: Pontine Central Gray
PCN: Paracentral nucleus of the thalamus
PDTg: Posterodrosal tegmental nucleus
PF: Parafascicular nucleus of the thalamus
PG (PN, BPN): Pontine gray
PGRN: Paragigantocellular reticular nucleus
PHY: Perihypoglossal nuclei
PMR: Paramedian reticular nucleus
PO: Posterior complex of the thalamus
PPN: Pedunculopontine nucleus
PPY: Parapyramidal nucleus
PRN: Pontine reticular nucleus
PRP: Prepositus nucleus
PRT: Pretectal region
PSV: Principal sensory nucleus of the trigeminal
RAmb: Midbrain raphe nucleus
RM: Nucleus raphe magnus
RN: Red nucleus
RPS: Relative Projection Strength
SAG: Nucleus sagulum
SC: Superior colliculus
SLC: Subceruleus nucleus
SLD: Sublaterodorsal nucleus
SNr: Substantia nigra, reticulata
SPIV: Spinal vestibular nucleus
SPVc: Spinal nucleus of the trigeminal, caudal
SPVi: Spinal nucleus of the trigeminal, interpolar
SPVo: Spinal nucleus of the trigeminal, oral
SUT: Supratrigeminal nucleus
SUV: Superior vestibular nucleus
TRN (NRTP): Tegmental reticular nucleus of the pons
V: Motor nucleus of the trigeminal
VAL: Ventral anterior-lateral complex of the thalamus
VEST: Vestibular nuclei
VII: Facial motor nucleus
VM: Ventromedial nucleus of the thalamus
VPL: Ventral posterolateral nucleus of the thalamus
VPM: Ventral posteromedial nucleus of the thalamus
VTA: Ventral tegmental area
X: Nucleus X
XII: Hypoglossal nucleus
Y: Nucleus Y
ZI: Zona incerta

## Acknowledgments

We thank Courtney Dobrott for sharing expertise in spinal cord removal, RNAscope methodology, and helpful comments on the manuscript. We thank Aya Miften for assistance in histology and Daniel Heck for work on preliminary datasets. We are grateful to Dr Jason Aoto for the preparation of AAVRetro viruses. This work was supported by NS114430, NSF CAREER 1749568, and by a grant from the Simons Foundation as part of the Simons-Emory International Consortium on Motor Control. Imaging of rabies label was performed in the Advanced Light Microscopy Core, part of the NeuroTechnology Center at University of Colorado Anschutz Medical Campus supported in part by Rocky Mountain Neurological Disorders Core Grant Number P30 NS048154 and by Diabetes Research Center Grant Number P30 DK116073 with the assistance of Dr. Radu Moldovan. Rabies viruses were obtained from the Salk GT3 Core Facility supported by NIH-NCI CCSG: P30 014195, an NINDS R24 Core Grant and funding from NEI.

---

**Supplemental Figure 1.**
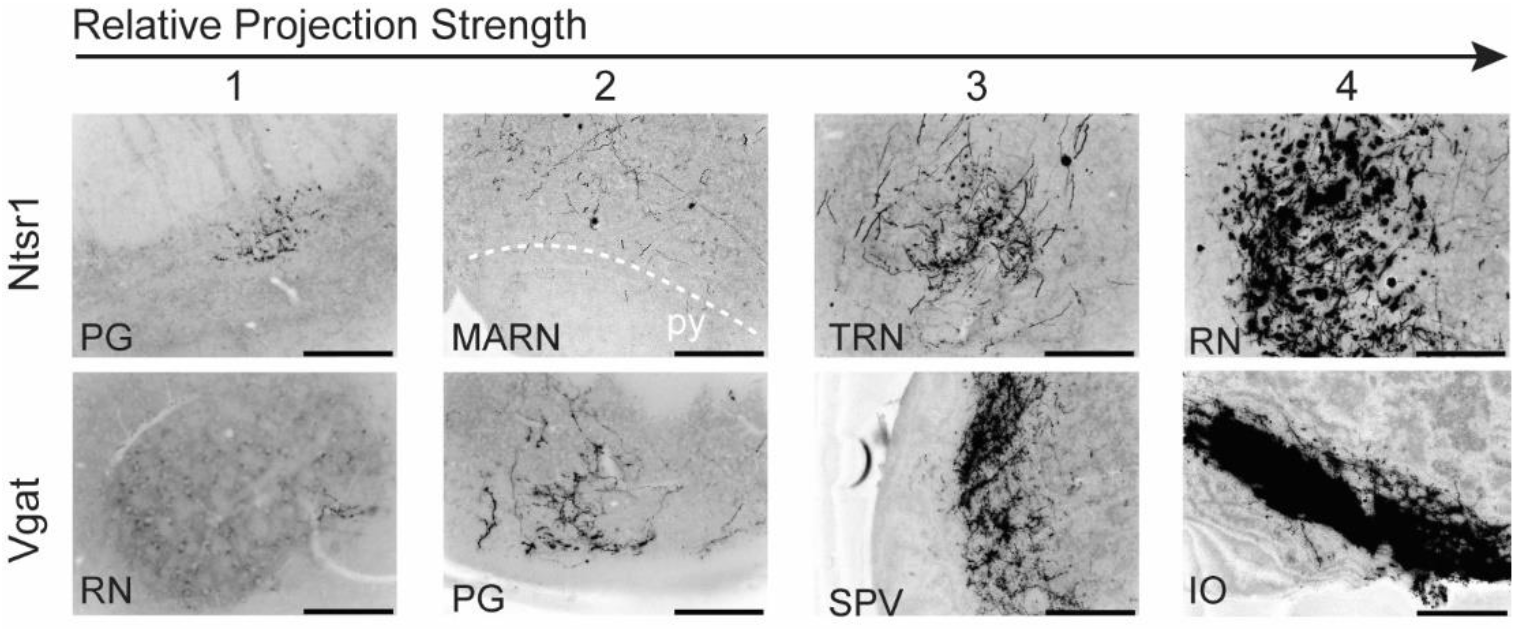
Example of semiquantitative scoring method of terminal field extent and density in Ntsr1 and Vgat-Cre mice. Top numbers indicate examples scoring ranging from 1 (sparse) to 4 (dense) innervation. *Pontine grey (PG), magnocellular reticular nucleus (MARN), tegmental reticular nucleus (TRN), red nucleus (RN), spinal trigeminal nucleus (SPV), inferior olive (IO)*.

**Supplemental Figure 2.**
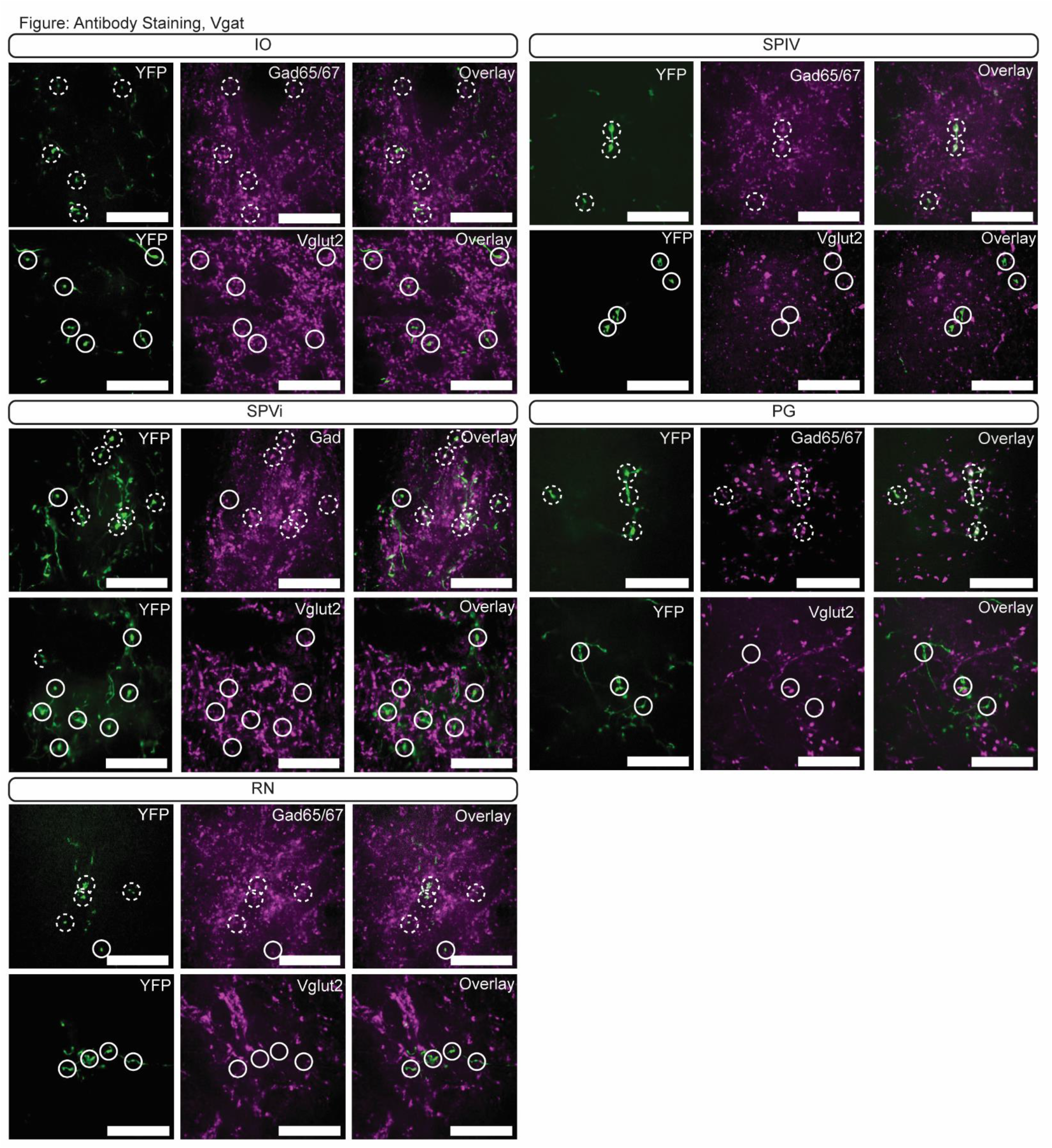
Immunoreactivity of Vgat-Cre terminal varicosities. YFP-positive terminals (green, left) in *inferior olive (IO), spinal vestibular nucleus (SPIV), spinal trigeminal nuclei, interpolar (SPVi), pontine grey (PG), and red nucleus (RN)* are stained for antibodies against Gad65/67 (top row; magenta; middle top) and Vglut2 (bottom row; magenta; middle bottom). Dashed circles indicate colocalized terminals while solid lines indicate a lack of colocalization observed in the two channels. Scale bars = 20 µms.

**Supplemental Figure 3.**
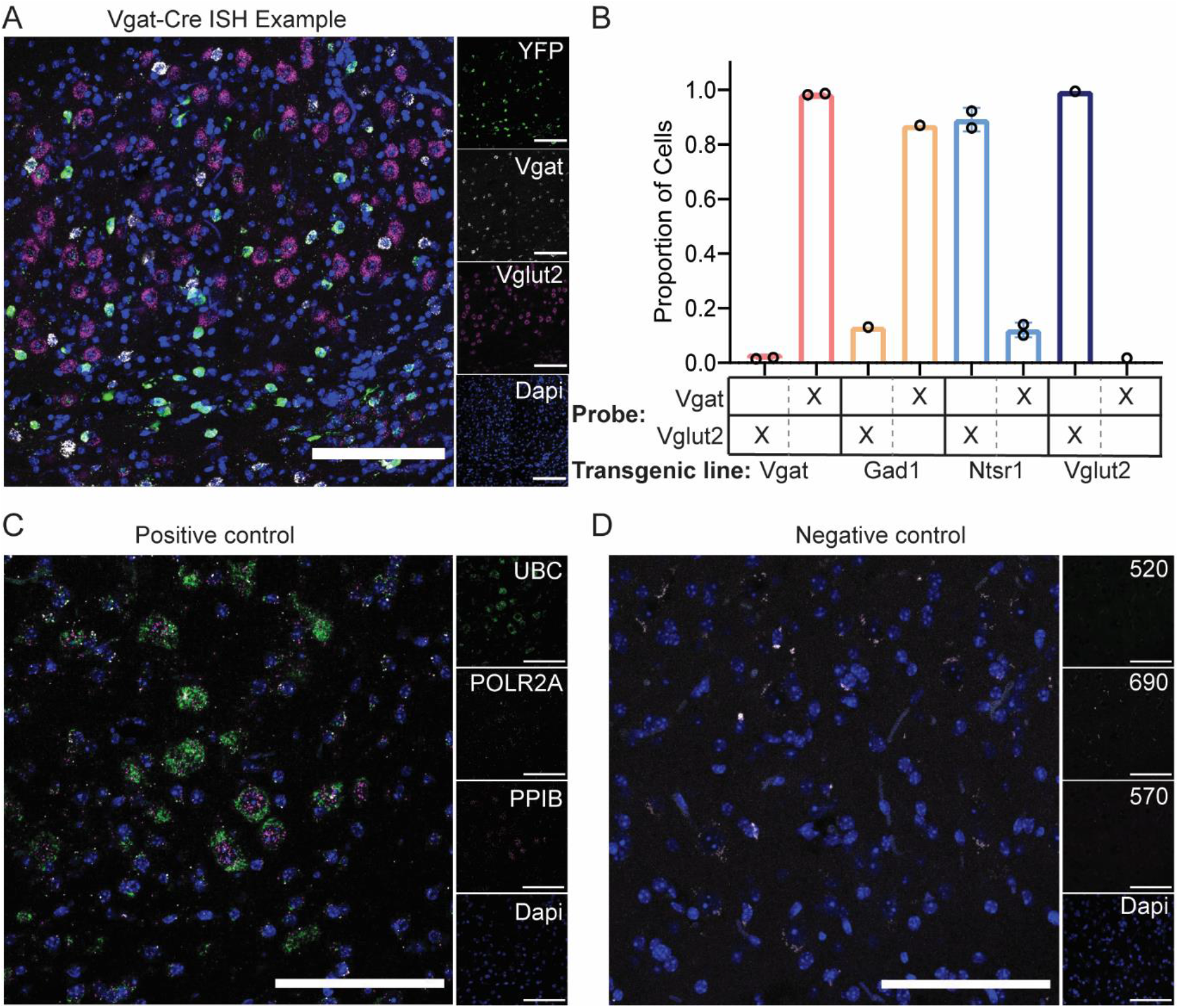
*In situ* hybridization methods and analysis. (A) Example image of *in situ* hybridization (ISH) images used for analysis. YFP-expressing cells (green; Opal 520) in a Vgat-Cre mouse co-localize with cells stained using an mRNA probe against SLC32a1 (Vgat; white; Opal 690), but do not co-localize with an mRNA probe against SLC17a6 (Vglut2; magenta; Opal 570). Cellular nuclei are stained with Dapi (blue). Scale bars = 200 µms. (B) Quantification of colocalization of YFP expressing cells in four transgenic mouse lines (Vgat-Cre, Gad1-Cre, Ntsr1-Cre, and Vglut2-Cre) with mRNA probes against Vgat an Vglut2. (C) RNAScope positive controls UBC (highest expressor; green; Opal 520 dye), POLR2A (lowest expressor; white; Opal 690 dye), PPIB (medium expressor; magenta; Opal 570 dye), and Dapi (blue) are shown to the right. Scale bars = 100 µms. (D) RNAScope negative controls (DapB; Opal 520 dye) with additional Opal dyes (570 and 690) incubated with tissue. Scale bars = 100 µms.

**Supplemental Figure 4.**
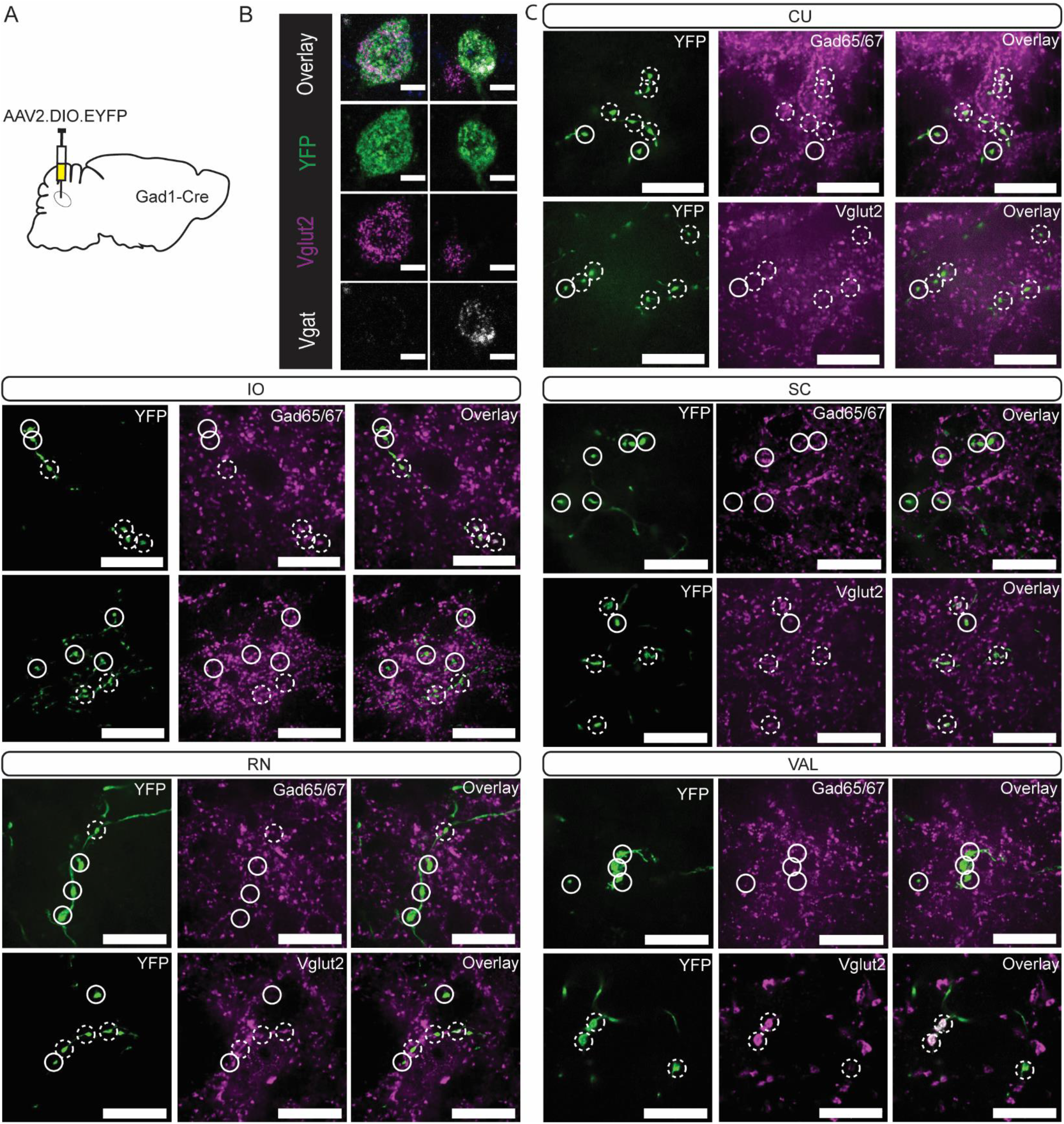
Gad1-Cre localized to mulitple cellular phenotypes in Int. (A) Schematic representation of experiment. (B) Example cells from *in situ* hybridization showing clear overlap with mRNA probes against SLC32a1 (Vgat) and SLC17a6 (Vglut2). Scale bars = 10 µms. (C) YFP-positive terminals (green) in *cuneate nucleus (CU), inferior olive (IO), superior colliculus (SC), red nucleus (RN), and ventral anterior-lateral complex of the thalamus (VAL)* are stained for antibodies against Gad65/67 (top; magenta) and Vglut2 (bottom; magenta). Dashed circles indicate colocalized terminals while solid lines indicate a lack of colocalization observed in the two channels. Scale bars = 20 µms.

**Supplemental Figure 5.**
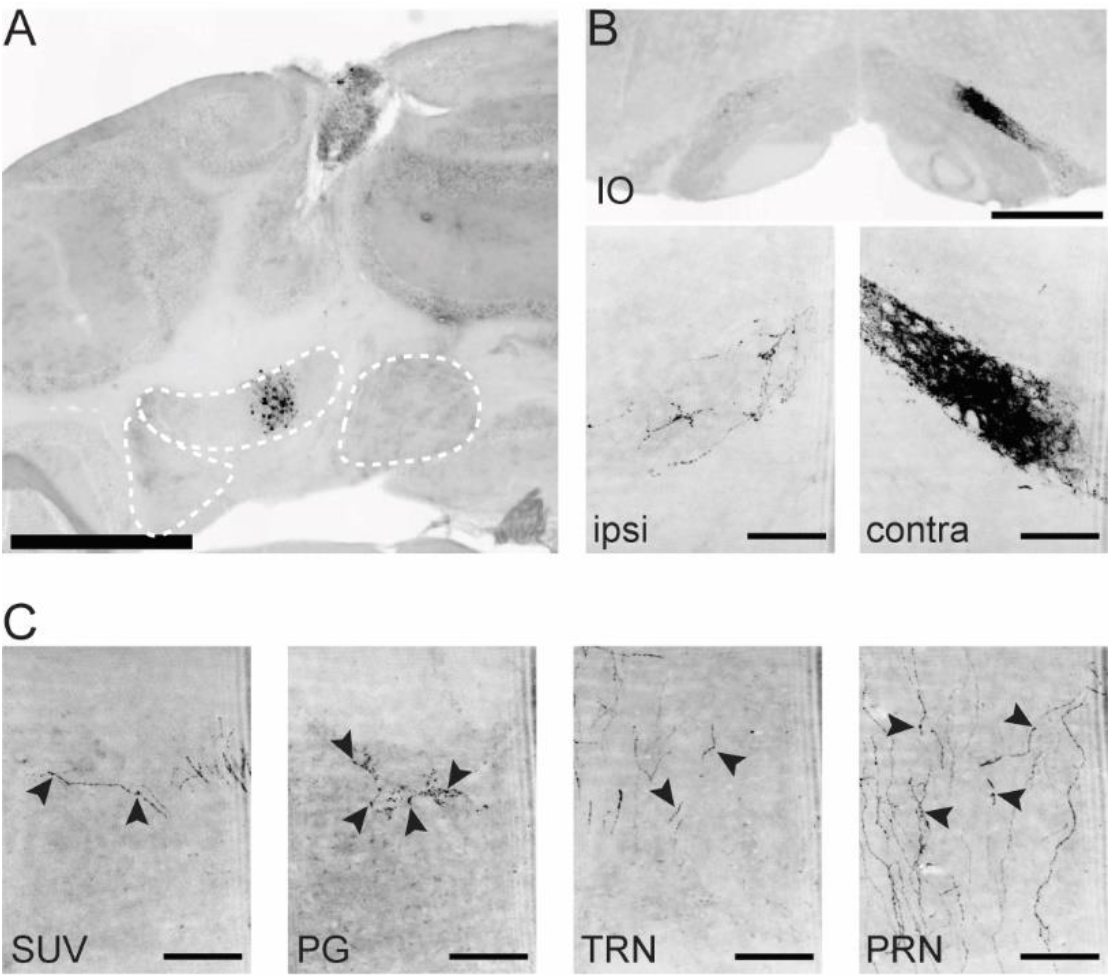
Example projections from an IntA restricted Int-Vgat specimen. (A) Example injection site. Scale bar = 1 mm. (B) Terminal contacts in ipsilateral (left) and contralateral (right) dorsal subnucleus of the *inferior olive (IO).* Scale bar = 500 µms (top) and 100 µms (bottom). (C) Terminal contacts in the s*uperior vestibular nucleus (SUV), pontine grey (PG), tegmental reticular nucleus (TRN), and pontine reticular nucleus (PRN).* Arrows denote example terminal varicosities on axonal processes. Scale bars = 100 µms.

**Supplemental figure 6.**
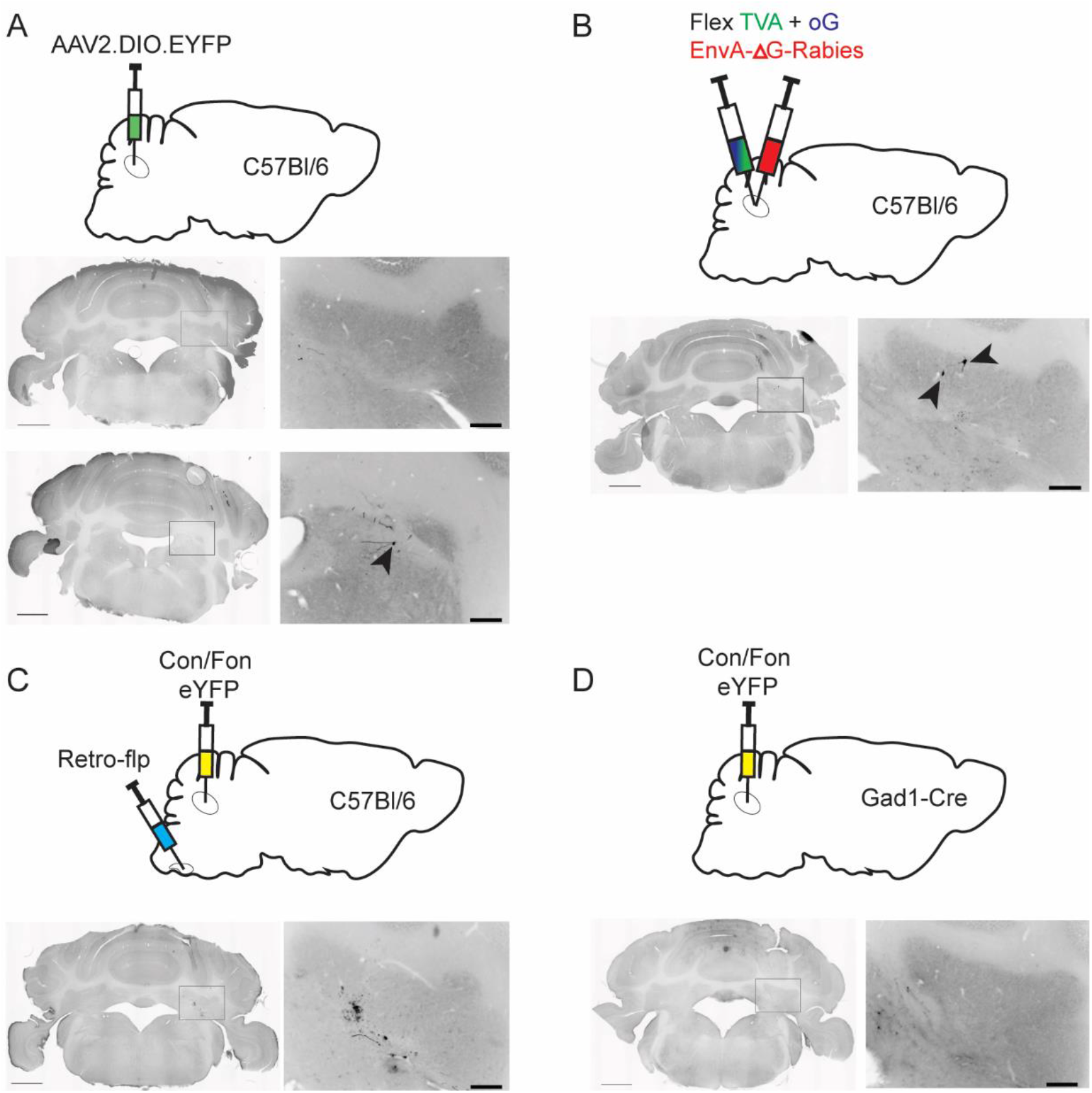
Viral control injections. (A) Example injection site (top) of AAV2.DIO.EYFP into a wildtype (C57Bl/6) mouse. Black arrowhead denotes singular fluorescent cell. (B) Example injection site of TVA, oG, and EnvA-ΔG-Rabies into a wildtype (C57Bl/6) mouse. Fewer than 10 cells were identified (denoted by black arrow heads) throughout the injection site location, but no retrogradely labeled cells outside of the cerebellar nuclei (CbN) were identified. (C) Example injection of Con/Fon-EYFP into Int of a wildtype (C57Bl/6) mouse after introduction of Retro.Flp to the contralateral *inferior olive (IO).* No cells were detected, though some tissue disruption/ autofluorescence was noted. (D) Example injection of Con/Fon-EYFP into a Gad1-Cre mouse. No cells were detected.

**Supplemental Figure 7.**
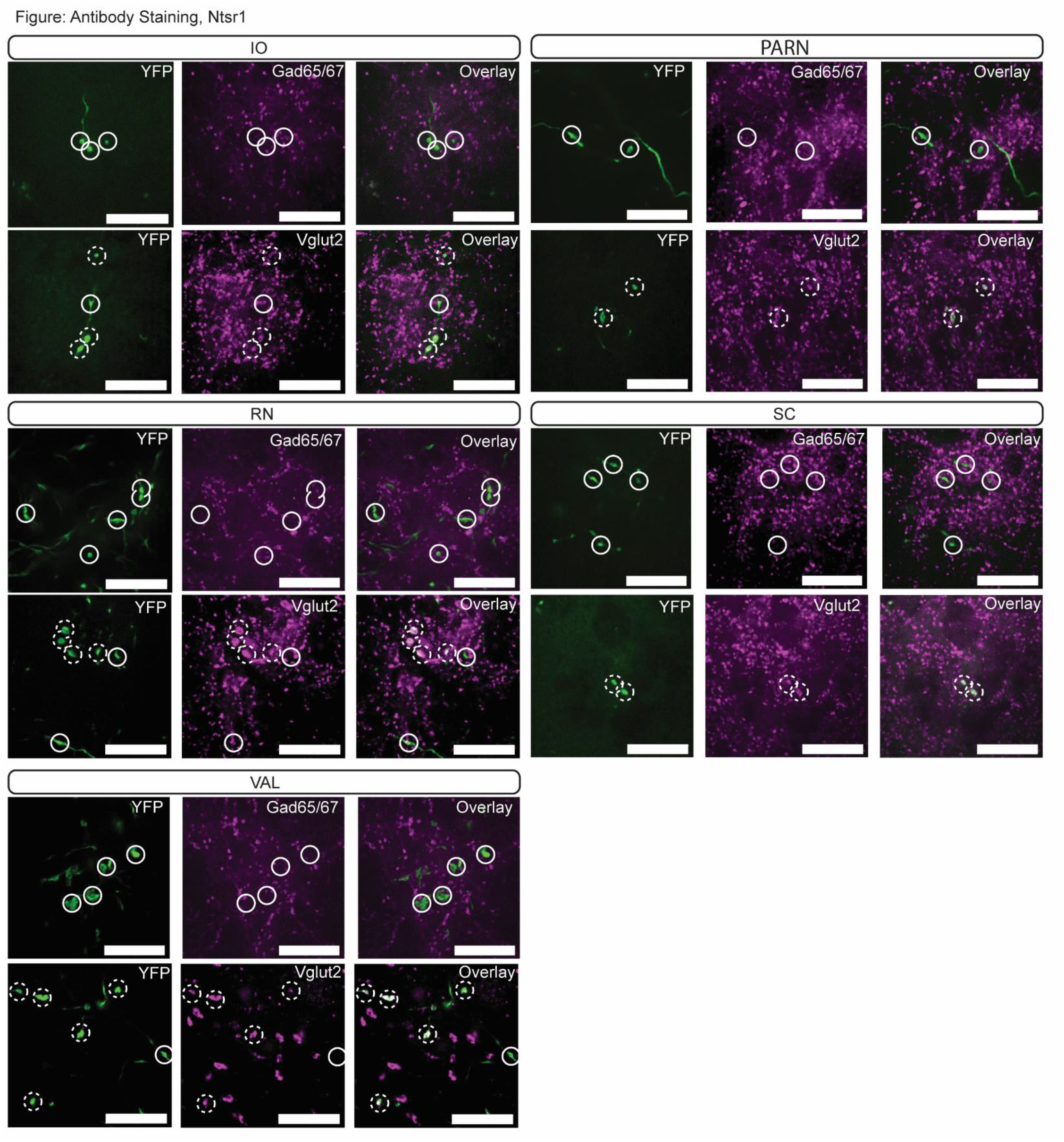
Immunoreactivity of Ntsr1-Cre terminal varicosities. YFP-positive terminals (green, left) in *inferior olive (IO), vestibular nuclei (VEST), spinal trigeminal nuclei (SPV), pontine grey (PG), and red nucleus (RN)* are stained for antibodies against Gad65/67 (top row; magenta; middle) and Vglut2 (bottom row; magenta; middle). Dashed circles indicate colocalized terminals while solid lines indicate a lack of colocalization observed in the two channels. Scale bars = 20 µms.

**Supplemental Figure 8.**
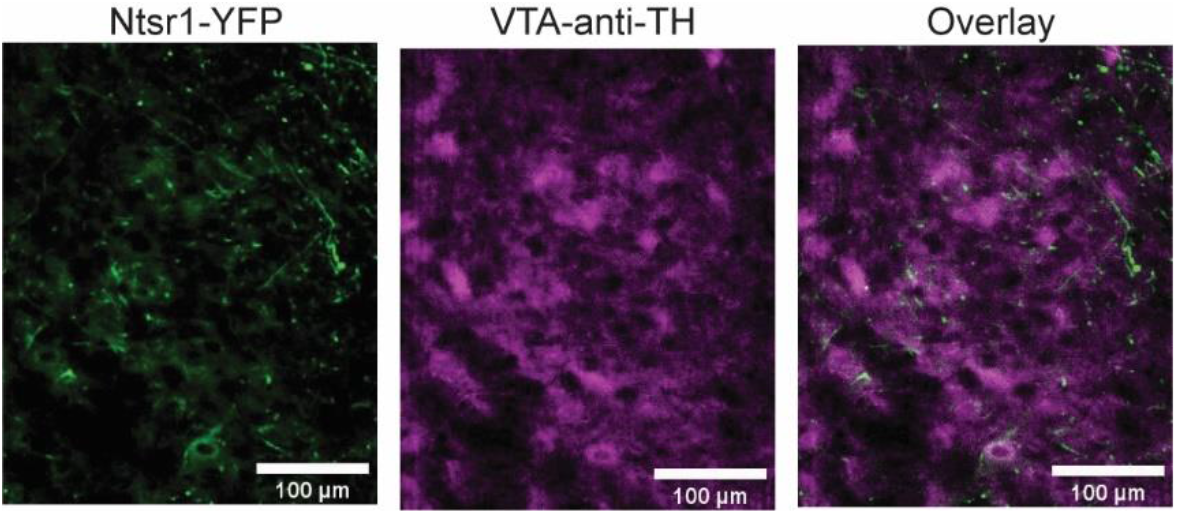
Int-Ntsr1 neurons (green) target tyrosine hydroxylase (TH; magenta) expressing neurons in the *ventral tegmental area (VTA)*. Scale bars = 100 µms.

**Supplemental figure 9.**
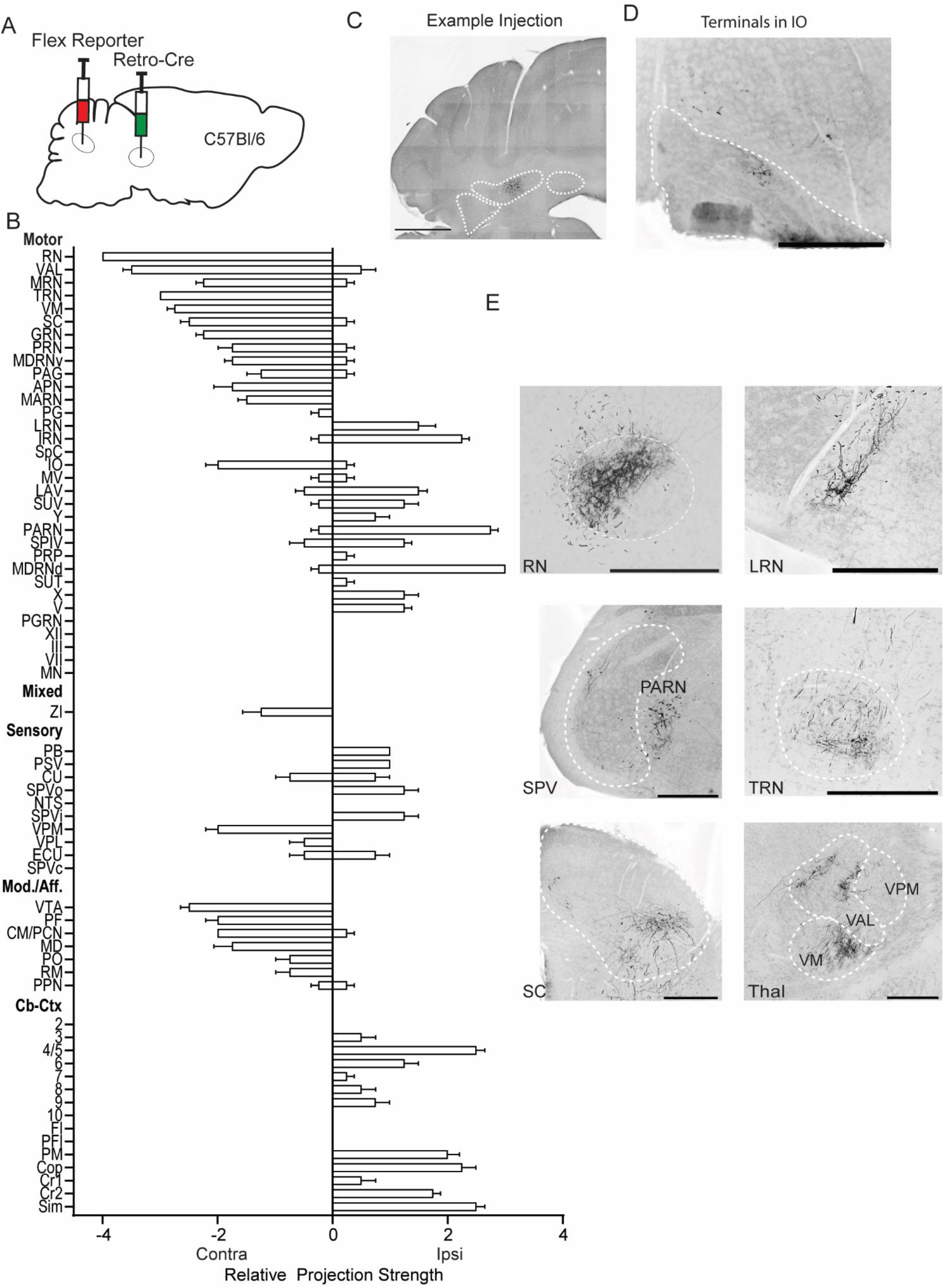
Labeling of RN-projecting Int neurons using viral Cre delivery. (A) Schematic of experiment. (B) Graphical representation of average projection strength in all targeted regions for Int-RN (n = 4). (C) Example injection site in wildtype C57Bl/6 mouse. The three main CbN are outlined in white (*Lateral Nucleus (LN), Interposed (IN), and Medial Nucleus (MN*) from left to right). Scale bar = 1 mm. Images are oriented as in Fig. 1. (D) Example terminal field in the *inferior olive (IO)*. (E) Example terminal fields within the *red nucleus (RN), lateral reticular nucleus (LRN), spinal trigeminal nucleus (SPV), parvicellular reticular nucleus (PARN), tegmental reticular nucleus (TRN), superior colliculus (SC), and several subdivisions of the thalamus (THAL).* Scale bars = 500 µms. *Ventral anterior-lateral complex of the thalamus (VAL), ventral medial nucleus of the thalamus (VM), and ventral posteromedial nucleus of the thalamus (VPM).\*

**Supplemental figure 10.**
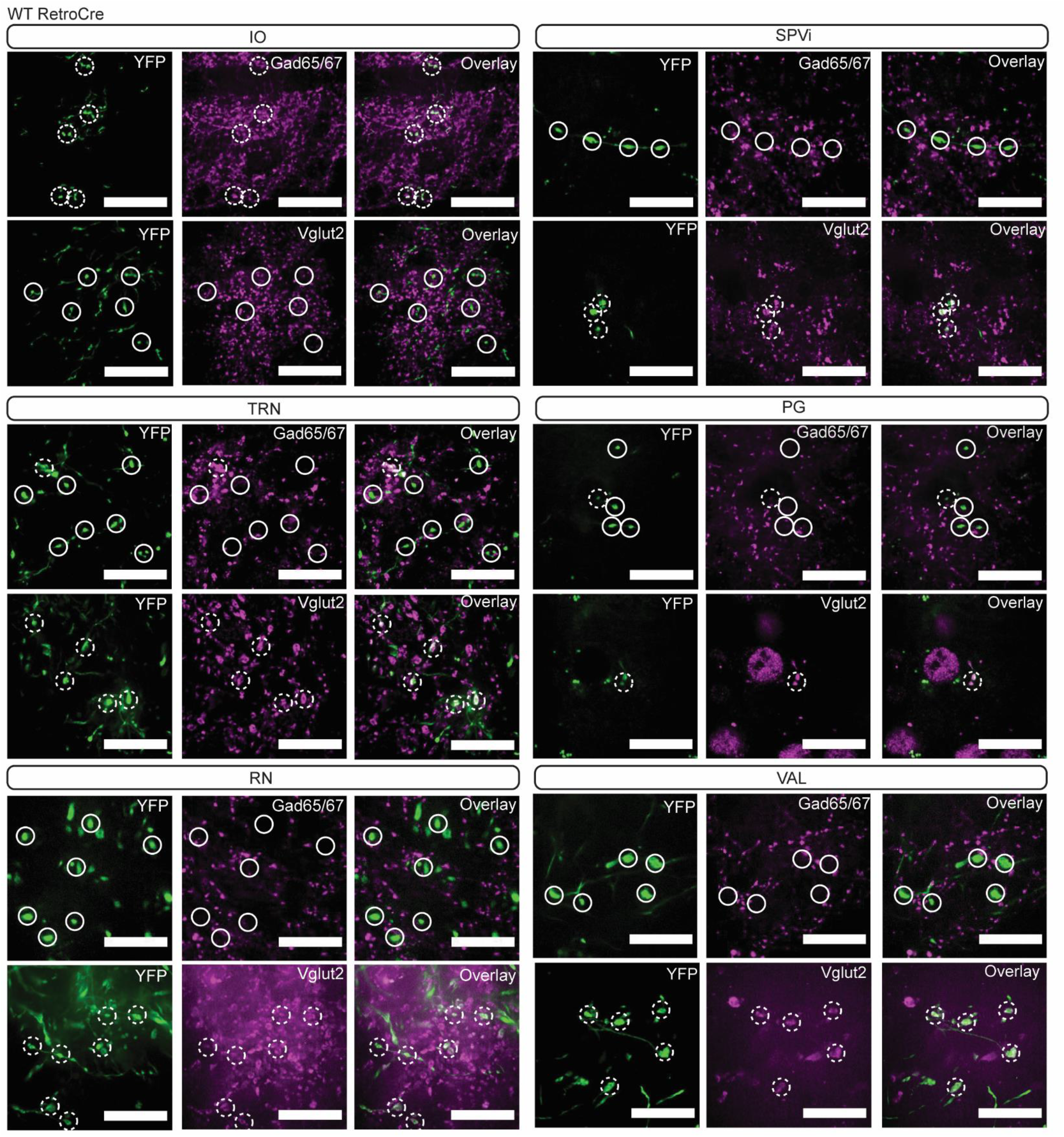
Immunoreactivity of Int-RetroCre^RN^ terminal varicosities. YFP-positive terminals (green, left) in *inferior olive (IO), spinal trigeminal nucleus, interpolar (SPVi), tegmental reticular nucleus (TRN), pontine grey (PG), red nucleus (RN), and ventral anterior-lateral complex of the thalamus (VAL)* are stained for antibodies against Gad65/67 (top row; magenta; middle) and Vglut2 (bottom row; magenta; middle). Dashed circles indicate colocalized terminals while solid lines indicate a lack of colocalization observed in the two channels. Scale bars = 20 µms.

**Supplemental Figure 11.**
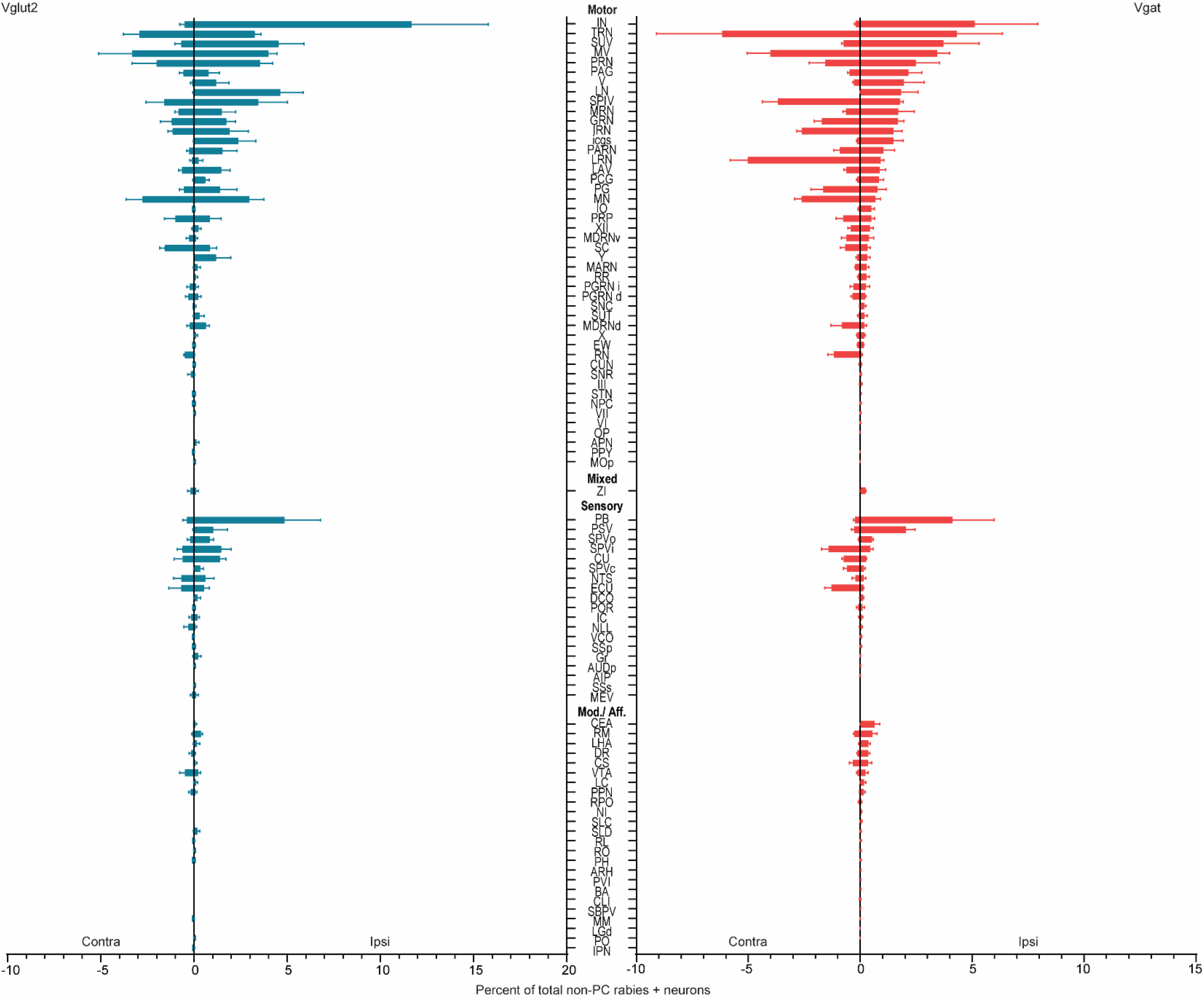
Summary of monosynaptically labeled inputs to Vglut2 (teal, n =3 mice) and Vgat (red, n =3 mice) neurons in IntA from extracerebellar regions. Mean and standard error are plotted. Note that the cerebellar nuclei included in the motor category did not express TVA and thus were not starter cells. See Table 1 for a full list of abbreviations.

